# Automated 3D Axonal Morphometry of White Matter

**DOI:** 10.1101/239228

**Authors:** Ali Abdollahzadeh, Ilya Belevich, Eija Jokitalo, Jussi Tohka, Alejandra Sierra

## Abstract

Axonal structure underlies white matter functionality and plays a major role in brain connectivity. The current literature on the axonal structure is based on the analysis of two-dimensional (2D) cross-sections, which, as we demonstrate, is precarious. To be able to quantify three-dimensional (3D) axonal morphology, we developed a novel pipeline, called ACSON (AutomatiC 3D Segmentation and morphometry Of axoNs), for automated 3D segmentation and morphometric analysis of the white matter ultrastructure. The automated pipeline eliminates the need for time-consuming manual segmentation of 3D datasets. ACSON segments myelin, myelinated and unmyelinated axons, mitochondria, cells and vacuoles, and analyzes the morphology of myelinated axons. We applied the pipeline to serial block-face scanning electron microscopy images of the corpus callosum of sham-operated (n = 2) and brain injured (n = 3) rats 5 months after the injury. The 3D morphometry showed that cross-sections of myelinated axons were elliptic rather than circular, and their diameter varied substantially along their longitudinal axis. It also showed a significant reduction in the myelinated axon diameter of the ipsilateral corpus callosum of rats 5 months after brain injury, indicating ongoing axonal alterations even at this chronic time-point.

## Introduction

Electron microscopy (EM) techniques are used extensively to assess brain tissue ultrastructure. Studies have reported the morphology, distribution, and interactions of different cellular components in both healthy and pathological brain using transmission electron microscopy (TEM)^1–4^. The ultra-thin sections prepared for TEM can only provide 2-dimensional (2D) information, limiting the full characterization of 3-dimensional (3D) cellular structures. Recent advanced EM techniques allow for new possibilities to study the ultrastructure of the brain in 3D^5–9^. One of these techniques is serial block-face scanning electron microscopy (SBEM)^6^. SBEM combines scanning electron microscopy (SEM) with back-scattered electron detection and low beam energies^10^. Images are acquired from the block-face of a sample each time an ultra-microtome inside the vacuum chamber removes the top section from a block-face to expose a new surface for imaging. The result is a stack of high-resolution, high-contrast images of tissue. Compared to other 3D EM techniques, such as focused ion beam (FIB), serial section TEM, or 3D-tomography, SBEM enables imaging of up to several hundreds of micrometers of tissue at nanoscopic resolution without manual tissue sectioning^5,11^. Thus, SBEM is the method of choice for mesoscale imaging of brain tissue ultrastructure.

Despite substantial progress in 3D image acquisition techniques, segmentation and quantification of SBEM data remain challenging. To date, several software tools have been developed that focus on either manual annotation (e.g., KNOSSOS^12^, TrakEM2^13^, Microscopy Image Browser^14^, and CATMAID^15^), or interactive processing of data by combining automated analysis and proof-reading capabilities (e.g., rhoANA^16^, ilastik^17^, and SegEM^18^). In addition to these software tools, a variety of studies have also proposed segmentation pipelines for analyzing large amounts of TEM data. Recent studies^19–26^ initially identified cellular boundaries using pixel-wise classification methods, followed by over-segmentation of the intracellular regions in each 2D image. This procedure requires merging the results within and between consecutive images using different strategies (e.g., watershed merge tree^23^, agglomerative or hierarchical clustering^19–21,25,26^, and joint segmentation of several images in anisotropic datasets^22,24^).

Although the EM segmentation approaches cited above have yielded impressive results, they have focused on the neuronal reconstruction of grey matter. In this study, we address quantification of white matter ultrastructure and particularly the morphometry of myelinated axons in sham-operated and animals after traumatic brain injury (TBI). Characterization of the white matter ultrastructure requires the segmentation of the white matter components from 3D-SBEM datasets. The previous segmentation methods cannot be used to address the segmentation of white matter for several reasons. First, using manual or semi-automated segmentation software tools (e.g.,^27^ TrackEM2^13^ and ilastik^17^) or pipelines (Chklovskii et αl.^20^), would be prohibitively time consuming. Providing annotated data, i.e., training data, for supervised learning-based segmentation methods, such as SegEM^18^ and SyConn^28^, is also very time consuming, or requires several annotators. Second, our interest lies in segmentation of several white matter constituents, such as myelin, myelinated and unmyelinated axons, cell bodies, mitochondria, and vacuoles. The methods utilizing binary pixel classifiers^23,24^, which assign a pixel as either a cell boundary or a cell-interior, are inappropriate for this type of multiclass segmentation problem. Especially if subcellular structures, such as mitochondria, are not labeled separately, the clustering step of these methods fails to correctly merge regions within a complete cell^26^. Some studies have addressed the multiclass segmentation^22,25,26^, and segmented mitochondria as a subcellular structure. These approaches, however, are only valid for grey matter, which does not contain myelin. In the SBEM images of white matter, the segmentation of mitochondria in the presence of myelin is extremely difficult because the signal intensity and textural features of mitochondria and myelin are highly similar. Interestingly, a previous study tracked axons in a SBEM volume of the optic tract using Kalman-snakes^29^ initialized either manually, or automatically using watershed filtering^30^. However, this approach fails in tracking full length of axons throughout the SBEM volume. Therefore, the automated segmentation of SBEM images of white matter requires a specifically developed method to address these problems. In conclusion, currently there exist no automated methods to quantify the axonal morphometry in 3D EM images.

We developed a novel pipeline for AutomatiC 3D Segmentation and morphometry Of axoNs (ACSON) in mesoscale SBEM volumes of white matter. The automated pipeline eliminates the need for time-consuming manual segmentation of 3D datasets and enables full 3D analysis of the white matter ultrastructure. ACSON segments the main cellular and subcellular components of the corpus callosum. To confirm the accuracy of the automated segmentation, the automated segmentation was evaluated against manual annotation by an expert. We quantified the actual cross-sections of the segmented myelinated axons based on their diameter and eccentricity. We analyzed the morphological features of SBEM datasets from the ipsilateral and contralateral sides of the corpus callosum in two sham-operated and three TBI rats.

## Results

### ACSON segmentation pipeline automatically annotates the white matter ultrastructure

We devised the ACSON segmentation pipeline to annotate the ultrastructure in SBEM volumes of white matter. The segmentation procedure of ACSON labeled white matter voxels as myelin, myelinated and unmyelinated axons, cells, mitochondria, and vacuoles. In addition, ACSON provided a separate label for each individual axon. The ACSON segmentation pipeline, illustrated in Fig. 1, comprised the following steps: 1) denoising the SBEM volume; 2) segmenting the volume using the bounded volume growing (BVG) technique, which integrates seeded region growing^31^ and edge detection algorithms in 3D; 3) refining the segmentation with supervoxel techniques; 4) segmenting the subcellular structure and cells, and annotating myelinated and unmyelinated axons.

**Figure 1.**
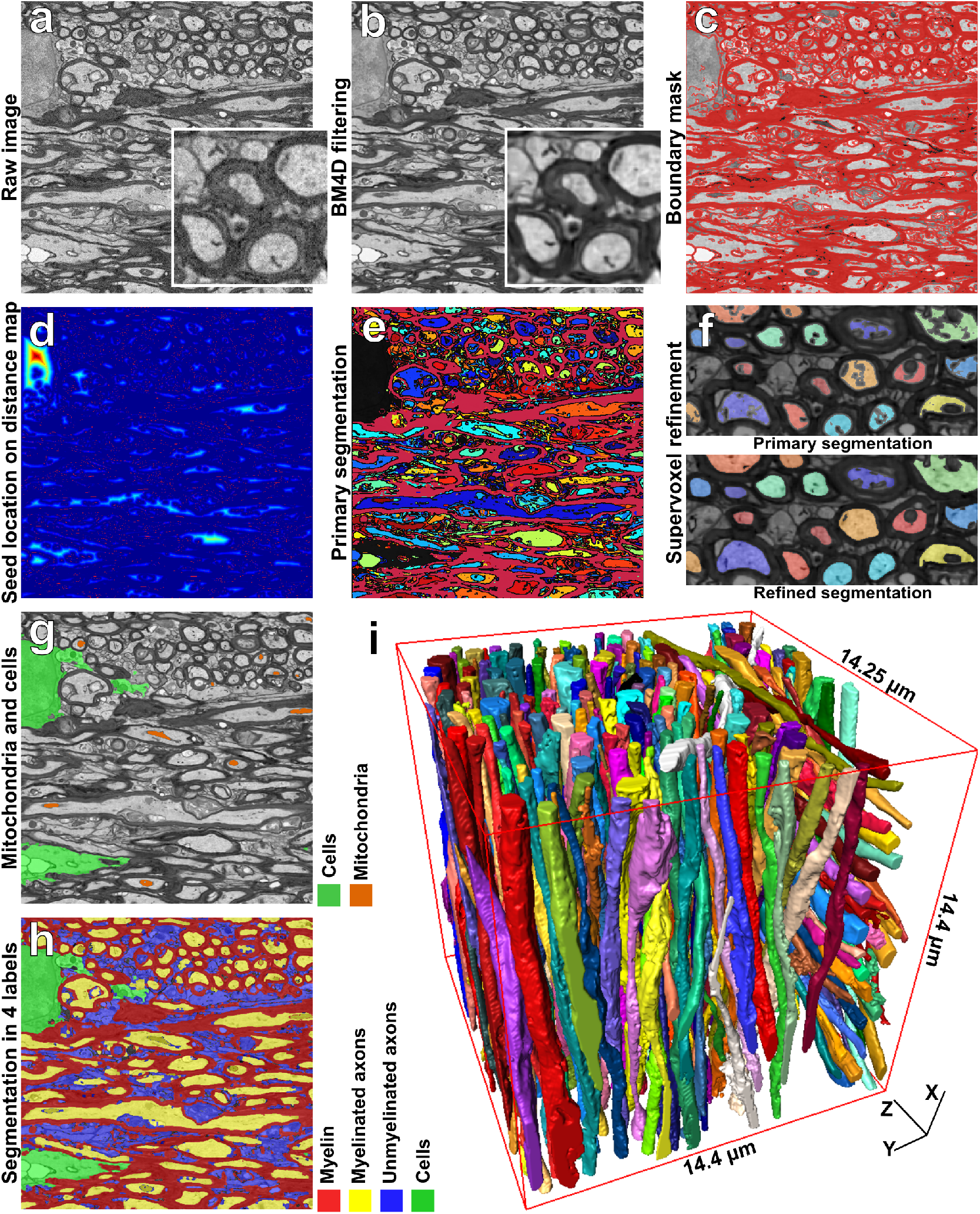
White matter ACSON segmentation pipeline. (**a**) A 2D representative image of the SBEM dataset from the contralateral corpus callosum of Sham-1 dataset. (**b**) The same image denoised with BM4D. (**c**) Boundary mask B: union of myelin obtained with BVG and dilated Canny edges. (**d**) The Euclidean distance transform was calculated for every slice of B individually to define the location of the seeds for the BVG algorithm. (**e**) The primary result of BVG segmentation. As the volume of cells/cell process exceeded ϑ, voxels corresponding to cells/cell process remained unlabeled in the primary segmentation (black regions). **(f)** The SLIC supervoxel technique refined the segmentation. **(g)** Segmentation of cells and mitochondria refined by the SLIC supervoxels. **(h)** A 4 labels map of myelin, intra-axonal space of myelinated and unmyelinated axons, oligodendrocyte cell body and its processes. **(i)** 3D rendering of myelinated axons in the contralateral corpus callosum of a Sham-1 dataset. The 3D rendering depicts myelinated axons with different thicknesses running along the dataset and organizing bundles with different orientations.

### Evaluation of the ACSON segmentation pipeline

We quantified the accuracy of the ACSON segmentation pipeline against manual segmentation by an expert. The expert (A.S.) manually segmented three 2D images (images 50, 55, and 60) from the contralateral dataset of one of the sham-operated rats (Sham-1). The images were selected to be 0.2 μm apart. The expert had no access to the automated segmentations of the dataset. Figure 2b shows the manual segmentations and the corresponding images produced by the automated segmentation. The segmentation accuracy was quantified using the precision (positive predictive value) and recall (sensitivity) in the tissue-type level similar to the previous studies^28,32^, and weighted Jaccard index and weighted Dice coefficients in the region level (see Materials and Methods). The precision and recall obtain their maximum value, one, if the automated segmentation correctly assigned voxels to myelin, myelinated or unmyelinated axon labels. The evaluation metrics in the tissue-type level, however, fail to account for topological differences between the ground truth and the automated segmentation. We used weighted Jaccard index and weighted Dice coefficients to evaluate the segmentation accuracy in the region level. Thus, each axon was considered to be its own region, which is a much more stringent evaluation criterion than considering all axons as a single region. The maximum value for these metrics is one, which occurs when a segmented region by ACSON perfectly matches a region segmented manually. If no overlap occurs, the metric is equal to zero. Table 1 reports the precision, recall, weighted Jaccard index, and weighted Dice coefficient values of the three slices shown in Fig. 2 and Supplementary Fig. S1. The results showed an excellent agreement between the automated and manual segmentations for myelin (precision ≥ 0.86, recall ≥ 0.88, weighted Jaccard index ≥ 0.78, and weighted Dice coefficients ≥ 0.87) and myelinated axons (precision ≥ 0.84, recall ≥ 0.88, weighted Jaccard index ≥ 0.80, and weighted Dice coefficients ≥ 0.88). For unmyelinated axons, in the tissue-type level the agreement was good (precision ≥ 0.72 and recall ≥ 0.68). The weighted Jaccard index and weighted Dice coefficients of unmyelinated axons showed approximately 0.37 and 0.50 agreement, respectively, which indicated topological differences and differences in perceiving ultrastructures such as myelin pockets and delamination between the automated and manual segmentations. These metrics are sensitive to minor displacements in the location of the boundaries as demonstrated in Supplementary Fig. S2.

**Figure 2.**
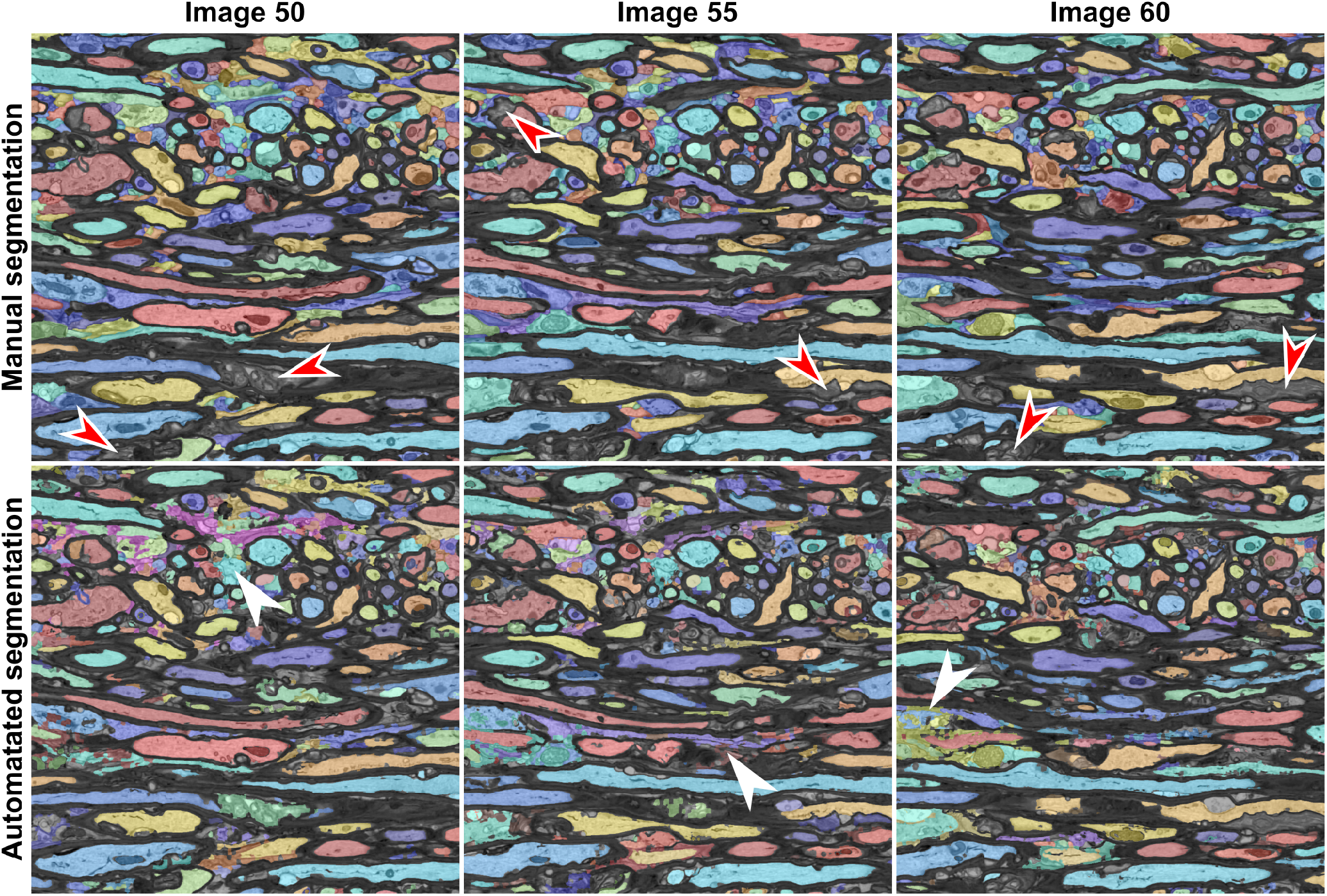
Manual expert segmentation and automated segmentation of three images from the contralateral corpus callosum of Sham-1 dataset. We used the Hungarian algorithm to match the color of segmented regions between automated and manual segmentation panels. The red arrowheads indicate delamination in the myelin sheaths. These substructures were annotated as myelin in the manual annotation, while the automated segmentation excluded the myelin delamination from the myelin-labeled structures. Occasionally, the automated segmentation erroneously merged neighboring unmyelinated axons when the space between the membranes was poorly resolved (white arrowheads). The automated segmentation of myelinated axons was in excellent agreement with the manual segmentation, as shown in Table 1.

**Table 1.** We evaluated ACSON segmentation by comparing it to the manual expert segmentation using precision (Prec.), recall (Rec.), weighted Jaccard index (WJI), and weighted Dice coefficient (WDC) metrics. The precision and recall metrics evaluate the segmentation in the tissue-type level and weighted Jaccard index and weighted Dice coefficients evaluate each axon as its own region. Therefore, the latter metrics are very sensitive to split-and-merge type errors.

|  | Image50 |  |  |  | Image 55 |  |  |  | Image 60 |  |  |  |
| --- | --- | --- | --- | --- | --- | --- | --- | --- | --- | --- | --- | --- |
|  | Prec. | Rec. | WJI | WDC | Prec. | Rec. | WJI | WDC | Prec. | Rec. | WJI | WDC |
| Myelin | 0.86 | 0.89 | 0.78 | 0.87 | 0.89 | 0.89 | 0.81 | 0.89 | 0.90 | 0.88 | 0.81 | 0.89 |
| Myelinated axons | 0.84 | 0.88 | 0.80 | 0.88 | 0.89 | 0.89 | 0.81 | 0.88 | 0.86 | 0.92 | 0.84 | 0.90 |
| Unmyelinated axons | 0.81 | 0.71 | 0.38 | 0.50 | 0.72 | 0.72 | 0.37 | 0.50 | 0.74 | 0.68 | 0.36 | 0.49 |

We further evaluated the segmentation of myelinated axons for the split (over-segmentation) and merge (under-segmentation) errors. Supplementary Fig. S3 illustrates the distribution of the length of myelinated axons in Sham-1 dataset. Myelinated axons with lengths smaller than 5 μm appeared in the corners of the SBEM volumes. These myelinated axons have partially traversed the SBEM volume. We did not observe split and merge errors in myelinated axons visually, confirming the excellent Jaccard index and Dice coefficient values for myelinated axons. The 3D rendering of myelinated axons for all datasets is shown in Supplementary Fig. S4. Also, Supplementary video shows the 3D rendering of myelin, cell, and myelinated axons in the contralateral corpus callosum of sham-1 dataset.

### ACSON morphometry pipeline automatically quantifies the segmented myelinated axons

The ACSON morphometry pipeline quantified cross-sections of the intra-axonal space of myelinated axons. A cross-section is the intersection of a segmented myelinated axon and a perpendicular plane to the axonal skeleton. To detect the axonal skeleton, we defined three points in the myelinated axon domain: one with the largest distance from the axon surface and two endpoints of the axons, i.e., the tips of the axon. The minimum-cost path connecting these three points was defined as the axonal skeleton (see Fig. 3a, which shows the cross-sectional planes at three randomly selected positions along an axon). The orientation of the cross-sectional planes at each skeleton point was determined by the unit tangent vector at that point. For each cross-section, we measured the length of minor and major axes of the fitted ellipses, equivalent diameter^33^ and eccentricity. The equivalent diameter is the diameter of a circle with the same area as the cross-section, and the eccentricity is a measurement of how much a conic section deviates from being circular. For example, the eccentricity of a circle is zero and the eccentricity of an ellipse is greater than zero but less than one. Figure 3b shows the measurements of minor axes along a myelinated axon. It is apparent that the myelinated axon diameter was not constant. We present more examples of cross-sectional variation along myelinated axons in Supplementary Fig. S5 and show that the histogram of cross-sectional diameters within a myelinated axon was bimodal, which can partially be related to the location of mitochondria^34^. Note that mitochondria was included in the intra-axonal space of myelinated axons, while vacuoles excluded, because they appeared between myelinated axon membrane and myelin.

**Figure 3.**
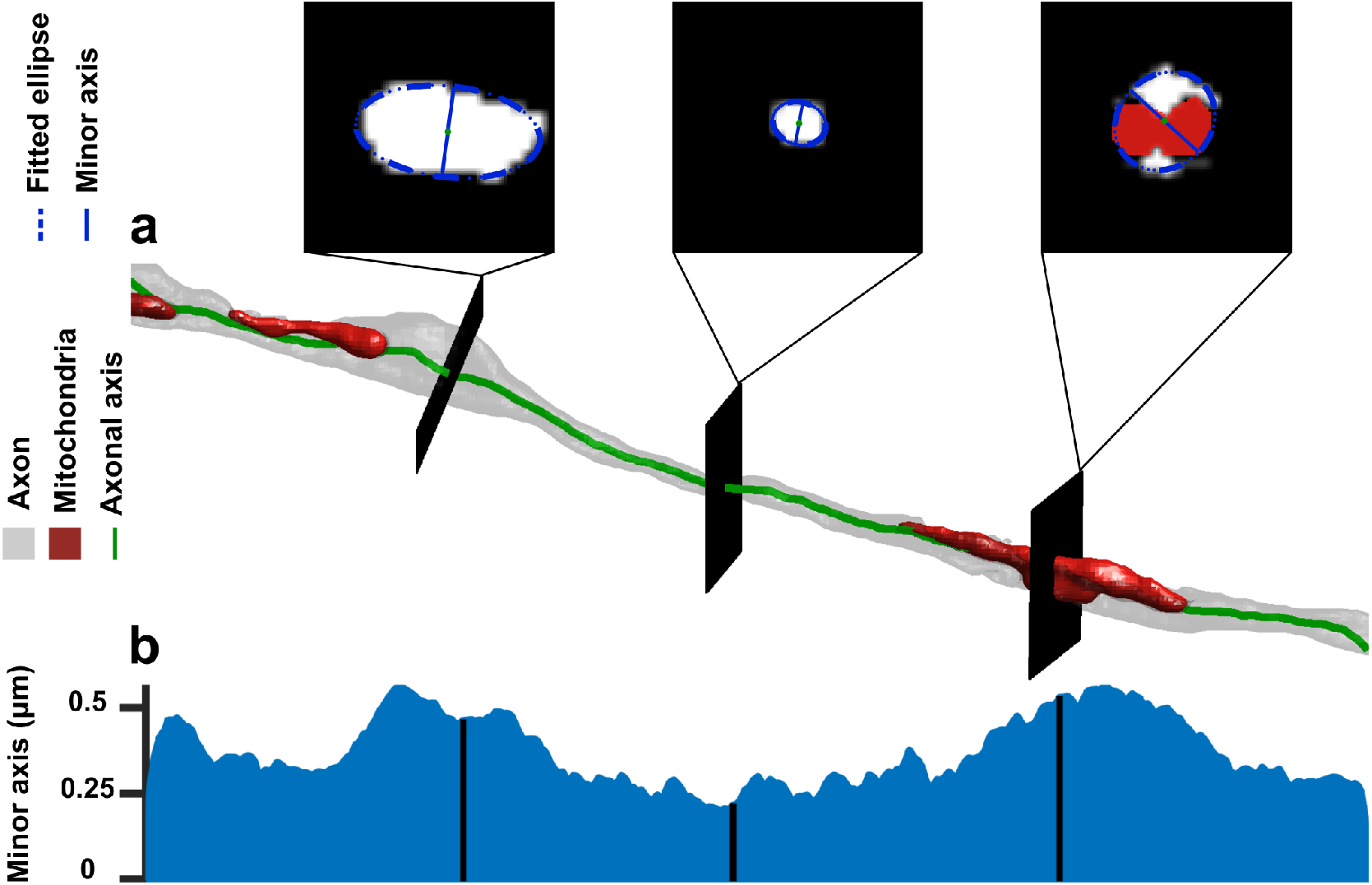
ACSON morphometry. (a) 3D reconstruction of a representative intra-axonal space of one myelinated axon and its mitochondria. Three intersecting planes at randomly selected positions show the cross-section of the myelinated axon. (b) The cross-sectional diameter of a myelinated axon varies along its length.

### Substantial differences between 2D and 3D morphological analyses

We compared the traditional 2D morphometry and the proposed 3D morphometry pipelines for myelinated axons in one dataset of the sham-operated rats (Sham-1). In Fig. 4a, a representative myelinated axon is elongated in the z direction. We sampled the myelinated axon parallel to the x-y plane at three points denoted as p1, p2, and p3 in the figure. Figure 4a shows that when the axonal axis was nearly perpendicular to the sampling plane (point p1), the relative difference between the 2D and 3D quantifications was small. However, when the axonal axis was not aligned with three main orientations, the relative difference between the 2D and 3D quantifications increased and was substantial (points p2 and p3). In addition, as the relative difference varied along a myelinated axon, a single 2D measurement was noisy. We compared the 2D and 3D measurements for all myelinated axons in contralateral corpus callosum of Sham-1 dataset as shown in Fig. 4b-e. The comparisons showed that the median of the relative difference, in percentage, was 16.64% for the minor axis, 18.25% for the major axis, 16.23% for the equivalent diameter, and 11.34% for the eccentricity. This indicates substantial differences between the 2D and 3D based measurements of morphometry of myelinated axons. As the cross-sectional measurements of a myelinated axon varied along it, sampling a single cross-section for measurement, as in 2D morphometry, gave an incomplete account of the parameters to be quantified. The 3D analysis, instead, quantified the morphology of a myelinated axon while taking into account the morphological variations along its axis.

**Figure 4.**
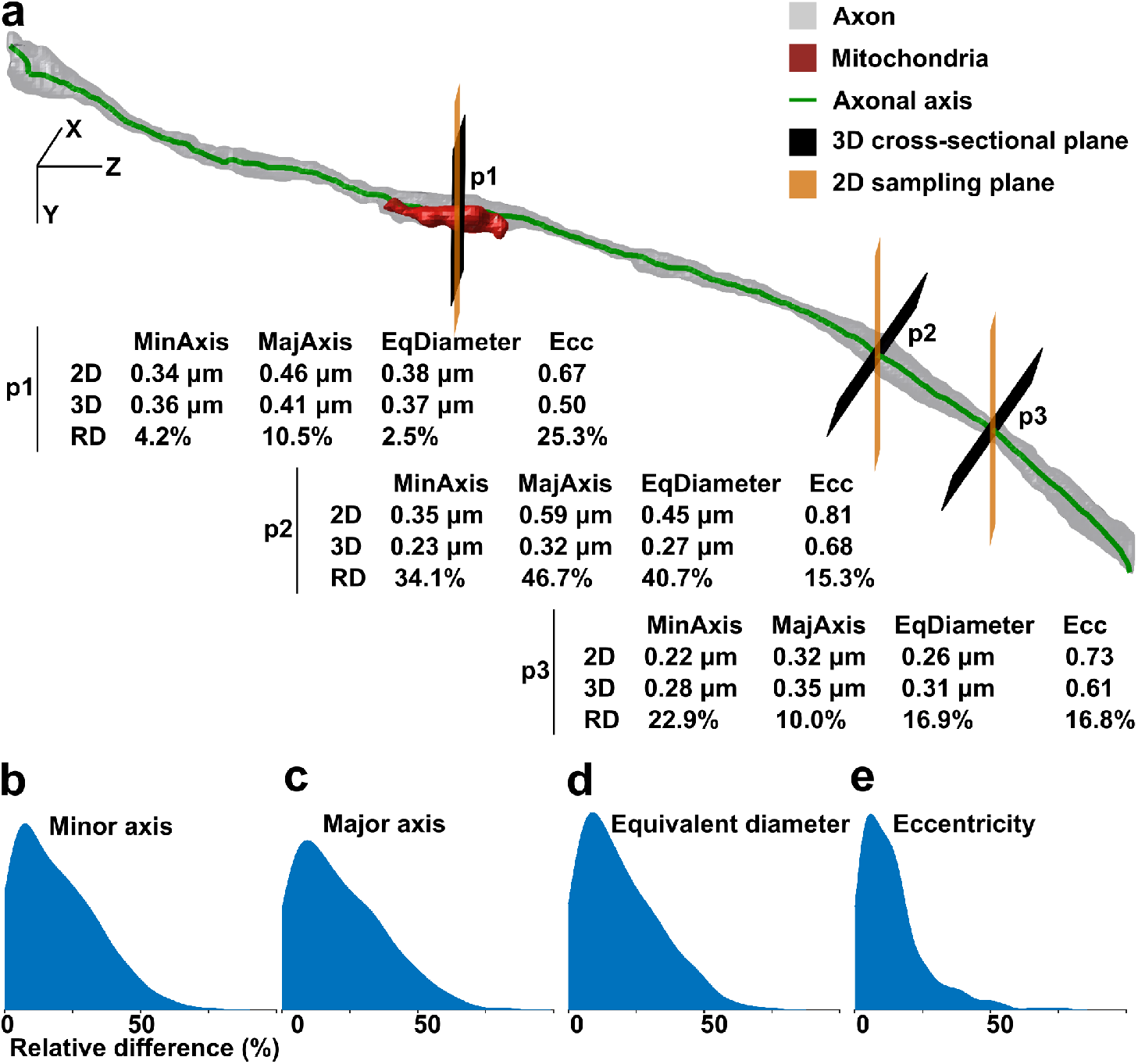
A comparison between the 2D and 3D morphological analyses. **(a)** 3D reconstruction of intra-axonal space of a representative myelinated axon with an axis that does not align with image cross-sections. The myelinated axon was quantified at three planes parallel to the x-y plane simulating the 2D morphometry and three cross-sections for the 3D morphometry. The quantified parameters were minor axis (MinAxis), major axis (MajAxis), equivalent diameter (EqDiameter), and eccentricity (Ecc). When the angle between the normal planes to the axonal axis and the image cross-sections grew, the relative difference (RD) between the 2D and 3D quantifications increased (p2 and p3). **(b-d)** Percentage of the relative difference between the 2D and 3D measurements of **(b)** the minor axis [median: 16.64 %, median absolute deviation (MAD): 10.66 %], **(c)** the major axis [median: 18.25 %, MAD: 16.68 %], **(d)** the equivalent diameter [median: 16.23 %, MAD: 9.60 %], and **(d)** the eccentricity [median: 11.34 %, MAD: 6.30 %] for all myelinated axons in contralateral corpus callosum of Sham-1 dataset.

### 3D morphometry of the ultrastructure of the corpus callosum

We quantified the morphological and volumetric aspects of the white matter ultrastructure in our SBEM datasets. For the morphological analysis, we thresholded myelinated axons based on their length, and preserved those myelinated axons whose length was long enough to run approximately one third of the SBEM volumes, which was equal to 5 μm. Supplementary Fig. S3a shows the thresholded myelinated axons which partially traversed the SBEM volume. In addition, the thresholding can eliminate myelinated axons which were split erroneously or subcellular structures that were mistakenly labeled as myelinated axons.

For each myelinated axon, we considered the median of the cross-sectional measurements (minor and major axes, equivalent diameter and eccentricity) as our measurement variable. The box plots of these measurements are shown in Fig. 5. We subjected these quantities to nested (hierarchical) 1-way analysis of variance (ANOVA) separately for each hemisphere^35^. The nested ANOVA tests whether there exists a significant variation in means among groups, or among subgroups within groups. The myelinated axons were nested under rats and the identities of rats were nested under the groups (sham-operated and TBI). We set the alpha-threshold defining the statistical significance as 0.05 for all analyses. The ANOVA was performed using the anovan function of MATLAB R2017b. Myelinated axon and rat identity were treated as random effects and the group was treated as fixed effect.

**Figure 5.**
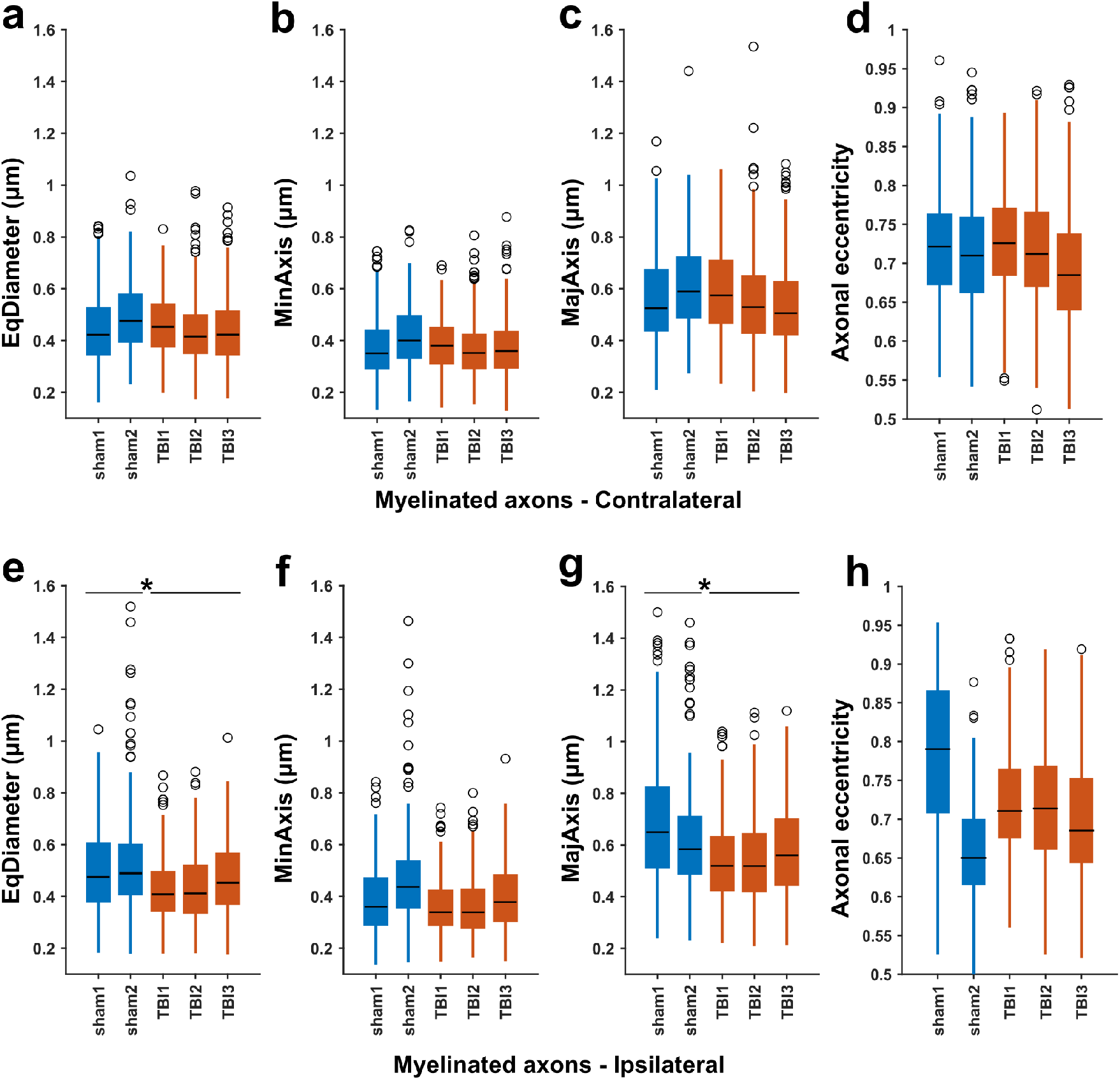
Box-plots of equivalent diameter (EqDiameter), minor axis (MinAxis), major axis (MajAxis), and axonal eccentricity measured for myelinated axons in sham-operated and TBI animals. The measurements are medians of the cross-sectional measurements of each myelinated axon. On each box, the central mark indicates the median, and the bottom and top edges of the box indicate the 25th and 75th percentiles, respectively. The whiskers extend to the most extreme data points not considered outliers, and the outliers are plotted individually using the ‘o’ symbol. Nested ANOVA showed a significant reduction in the diameter of myelinated axons (the equivalent diameter and the length of major axis) in the ipsilateral corpus callosum of rats after TBI.

We observed a significant reduction of the equivalent diameter (F = 14.4, p = 0.029) and the major axis (F = 26.4, p = 0.012) in the ipsilateral corpus callosum of TBI rats as compared to the ipsilateral corpus callosum of sham-operated rats (see Supplementary Table S1 and Fig. 5). The contralateral corpus callosum, however, did not show a significant difference between sham-operated and TBI rats for any of the measures (see Supplementary Table S1 and Fig. 5). We measured that the equivalent diameter was about 15 % greater than the minor axis of the fitted ellipses. The eccentricity in all datasets were markedly different from zero indicating that the axonal cross-sections were not circular, but elliptical.

We also partitioned the variability of the measurements in cross-sectional, axonal, and animal levels, known as variance component analysis. The variance components describe what percentage of the total variance is attributable to each level. The design matrix of a 4-levels nested ANOVA for all measurements was very large (2 groups, i.e., sham-operated and TBI, 5 animals, approximately 250 myelinated axons per animal, and approximately 250 cross-sectional measurements per myelinated axon) making it impossible to perform the computations with the complete dataset. Therefore, we sampled 10000 of the measurements (without repetition), computed the variance components and repeated the procedure for 10 times. The variance components (Table 2) indicated that the greatest part of the variance was attributed to variance of cross-sections, then between myelinated axons, and the least amount of variance was attributed to between sham-operated and TBI animals.

**Table 2.** Variance components analysis. The variance was partitioned into cross-sectional, axonal, and animal levels.

|  |  | Cross-sections (%) | Axons (%) | Animals (%) |
| --- | --- | --- | --- | --- |
| Contralateral | EqDiameter | 57.33 ± 1.09 | 41.29 ± 1.16 | 1.38 ± 0.46 |
|  | MinAxis | 59.47 ± 0.93 | 39.14 ± 1.00 | 1.39 ± 0.30 |
|  | MajAxis | 63.59 ± 1.18 | 35.15 ± 1.09 | 1.26 ± 0.31 |
|  | Eccentricity | 83.12 ± 0.70 | 15.97 ± 0.66 | 0.91 ± 0.20 |
| Ipsilateral | EqDiameter | 49.85 ± 1.26 | 48.02 ± 1.26 | 2.13 ± 0.46 |
|  | MinAxis | 51.67 ± 1.12 | 44.47 ± 1.12 | 3.86 ± 0.30 |
|  | MajAxis | 56.18 ± 1.08 | 41.27 ± 1.23 | 2.55 ± 0.36 |
|  | Eccentricity | 75.53 ± 0.98 | 17.96 ± 0.86 | 6.51 ± 0.51 |

We also quantified the volumetric aspects of myelin in our 3D-SBEM datasets. The ultrastructure volumetry was dataset-dependent, preventing a direct comparison between datasets. For example, the volume of cell body/processes varied among datasets [1.3% − 22.1%], influencing the volume of the other ultrastructures (see the results of volumetric analysis in Supplementary Table S2). Therefore, we calculated the implicit representation of the g-ratio^36^ with no measurement of the myelin thickness^37^, denoted as *aggregate* 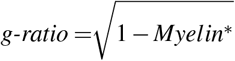. We defined *Myelin*\* as the ratio of the myelin volume to the myelin volume plus the intra-axonal volume of all myelinated axons. In addition, we calculated the density for myelinated axons and mitochondria defined as 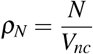, where *N* is the number of myelinated axons or mitochondria counted once in a volume *V_nc_* which is the volume of non-cellular ultrastructure. Supplementary Table S2 shows the aggregate g-ratio and the density of myelinated axons and mitochondria for all datasets. We did not run statistical hypothesis tests for comparing these metrics between sham-operated and TBI rats. However, visually, we observed myelin delamination and pockets in myelin sheath more frequent in rats after TBI compared to sham-operated rats.

### Computation time

On a 4-core Intel CPU 3.41 GHz machine with 64 GiB RAM using MATLAB R2017b, the computation times for one dataset were approximately as follows: block-matching and 4D (BM4D) filtering 5-7 h, segmentation process 1 day, and skeletonization and cross-sectional analysis 6 h. Correcting the segmentation of mitochondria and proofreading of myelinated axons versus unmyelinated axons was accomplished in 7 h for the entire SBEM-volume. The manual expert annotation of a single slice of the SBEM datasets required 10 h.

## Discussion

Previous studies that quantified white matter were limited to 2D morphometry, which simplifies the assumptions about axonal morphology. In this paper, we reported an extensive 3D morphological analysis of SBEM volumes. For this, we devised a novel pipeline, termed ACSON, for automated segmentation and morphometry of the cellular and subcellular components of the corpus callosum in SBEM datasets. ACSON segmented white matter into myelin, the intra-axonal space of myelinated and unmyelinated axons, cell bodies and their processes, and subcellular compartments, such as mitochondria and vacuoles. The segmentation accuracy evaluations revealed excellent agreement between the automated and manual segmentation of myelin and myelinated axons. ACSON quantified the morphology of the segmented myelinated axons. The 3D morphometry demonstrated a substantial variation in the diameter of myelinated axons along their longitudinal axis. The results indicated that the cross-sections of a myelinated axon are more likely to be elliptical than circular. To compare sham-operated and TBI animals, we used nested ANOVA and variance component analysis. After TBI, we found a significant reduction in the diameter of myelinated axons in the ipsilateral of corpus callosum, indicating that the alterations persisted for several months after the injury.

Traditionally, studies have modeled axons as straight cylinders, and utilized 2D-EM sections to assess the axonal morphology^38–40^. However, studies of unmyelinated axons in peripheral nerves^41^ and in the hippocampus and cerebellum^42^ indicated highly irregular axial shapes with periodic varicosities containing organelles^42^. In addition to studying the anatomical alterations of brain ultrastructure in disease, models of diffusion magnetic resonance imaging (dMRI^)43,44^ or electrophysiology^45,46^ in many studies assume simplified geometries for axons. However, Novikov *et al*.^47^ and Fieremans *et al*.^48^ have shown the effect of structural disorder along axons and micron-level sample architecture while modeling dMRI. Thus, realistic modeling of cells^49^ and myelinated axons^50^ have raised attention for numerical simulation of dMRI. Segmentation of high-resolution EM volumes provides realistic properties such as axonal diameter, eccentricity of cross-sections and orientation dispersion that can substitute simplified models of axons.

Our segmentation and morphometry pipeline is automated. The ACSON segmentation pipeline requires tuning several parameters, such as the thresholds for measuring similarity or vacuole intensity. These parameters, however, are easy to set. Annotating the mitochondria was the only step that remained a challenge and required human intervention. Compared with manual segmentation, which required, on average, 10 hours for a single slice, annotating the mitochondria required approximately six hours for the entire dataset comprising 285 SBEM images. In an earlier study, Lucchi *et al*.^32^ specifically targeted segmentation of mitochondria in high resolution FIB-SEM datasets of 5-6 nm x 5-6 nm planar resolution, which is a much finer resolution than in our datasets. They reported that mitochondria and myelin were difficult to discriminate whenever they were in close proximity. The high resolution of their FIB-SEM datasets enabled them to outline the prominent shape-features of mitochondria and they segmented mitochondria almost in the absence of myelin. Unfortunately, their technique is not applicable to our data due to our larger tissue samples, lower resolution, and proximity of mitochondria to myelin.

The ACSON morphometry pipeline required no user input parameters and extracted a sub-voxel precise and naturally smooth axis for each individual myelinated axon. We assumed a myelinated axon skeleton with no branches and only two endpoints. This allowed us to optimize the computation time for skeleton extraction. The computational efficiency was crucial because we solved the eikonal equation for several thousands of axons with multi-stencils fast marching (MSFM), which is more accurate but also more time-consuming than the fast marching method (FMM)^51^. We quantified myelinated axons for their minor and major axes, equivalent diameter and eccentricity. The minor and major axes and the equivalent diameter, all can describe the cross-sectional diameter of myelinated axons. Estimating the minor and major axes, however, requires fitting an ellipse to the cross-sections. Ellipse fitting step regularizes the measurements, which reduces the cross-sectional variance along myelinated axons. The equivalent diameter is measured directly from the area of cross-sections and it can capture the cross-sectional variance along myelinated axons, however, it measured the cross-sectional diameter 15% greater than the minor axis. Our morphological analyses of myelinated axons in sham-operated animals were in line with a previous study^52^ measuring axon diameter in the rat corpus callosum. Kim *et al*.^52^ measured an average myelinated axon diameter of 0. 35 μm in the splenium of the corpus callosum in male and female rats at postnatal day 60 using 2D electron microscope. The authors measured the myelinated axon diameter as the minimum diameter of each axon. For a comparison to this study, we measured an average myelinated axon diameter as equivalent diameter (Sham-1: 0.44 μm and Sham-2: 0.5 μm) and minor axis (Sham-1: 0.37 μm and Sham-2: 0.42 μm) for the contralateral corpus callosum. Note that our tissue samples were acquired from the body of the corpus callosum at approximately −3.80 mm from bregma from male adult rats. Moreover, the variability of the myelinated axon diameter along its longitudinal axis can be related to the accumulation of organelles, such as mitochondria, increasing the cross-sectional diameter locally^34^. Wake *et al*.^53^ also indicated that accumulation of vesicles containing neurotransmitter along an axon induces a local rise in the form of varicosities.

The segmentation accuracy evaluations demonstrated substantial agreement between the automated and manual segmentations as shown by the precision and recall metrics. These evaluations, however, did not provide a realistic estimation of the segmentation quality when the goal is to separate distinct axons. The weighted Jaccard index and weighted Dice coefficients computed in the region level were much more stringent evaluation measurements, and demonstrated excellent results for the segmentation of myelin and myelinated axons. However, the weighted Jaccard index and weighted Dice coefficients indicated 37% and 50% agreement in the segmentation of unmyelinated axons, respectively. There were several possible reasons for the decreased accuracy in the segmentation of unmyelinated axons. First, pockets in the myelin sheaths were included in the myelin label, while ACSON annotated these volumes individually. This potentially introduced false positives reducing the weighted mean of the Dice coefficients. Second, faintly resolved unmyelinated axons might have been included into the cell body/process annotation. Finally, the cellular boundaries of unmyelinated axons were often difficult to detect, which resulted in the erroneous merging of neighboring unmyelinated axons. Overall, as the precision and recall scores of unmyelinated axons were good, we investigated the volumetry of unmyelinated axons.

We have provided a framework for the statistical inference of data with nested structure. Simply pooling the measurements from all animals is not a valid approach^35^. Also, averaging the lowest levels of hierarchy has sub-optimal statistical power^54^. The nested ANOVA analysis, however, can capture the variability of the measurements in all levels. In addition to testing the equality of the means at each level through nested ANOVA, we partitioned the variance into different levels. The variance component analysis showed that variation among animals was relatively small compared to the variation among cross-sections and myelinated axons.

It is becoming increasingly clear that white matter pathology plays a major role in brain disorders. After TBI, the white matter pathology is extremely complex. The initial response of an axon to a brain injury^55^ can be either degeneration or regeneration. Studies of the white matter ultrastructure indicate axonal swelling in the early stages of TBI^56,57^. Axonal damage persists for years after injury in humans^58^ and for up to 1 year in rats^59^. In the present study, we observed morphological changes in the corpus callosum 5 months after severe TBI in rats. We found that the axonal diameter of myelinated axons in the ipsilateral corpus callosum of TBI rats had reduced significantly. The reduction of the diameter of myelinated axons might indicate a prolonged axonal degeneration^60^ or regeneration^55^. In analysis of myelin, the automated pipeline has excluded the pockets and delamination from the myelin label. Visually, we observed more frequent delamination in TBI animals, which might indicate active myelin processes still ongoing 5 months after the injury. Myelin delamination was also observed in sham-operated rats, which may be part of the natural dynamics of healthy myelin. Note that, we observed pockets in the myelin sheaths more frequently in animals after injury. The delamination of the myelin sheaths in healthy and injured brains at this chronic time point is currently unclear, and warrants further studies.

When characterizing the ultrastructural morphometry, the tissue shrinkage caused by fixation, staining, and sectioning might affect quantification of the axonal diameter and volumes^61^. In addition, the locations from which the specimens were obtained might influence the quantifications. The SBEM datasets in this study were consistently imaged at a specific location in the corpus callosum in both sham-operated and TBI animals, and in both hemispheres. Due to the small tissue size, however, the environment might change from one sample to another. For example, one of our datasets from the contralateral corpus callosum in a TBI rat contained more cell body/process volume than the other datasets. The small sample size can be reflected as well in the orientation of the myelinated axons. In a coronal view of the rat brain, the corpus callosum mainly contains in-plane parallel axons oriented in the latero-medial orientation, however, there are also bundles of axons oriented dorso-ventrally, and even rostro-caudally. For that reason, our datasets can contain different axonal populations with different orientations. Thus, studies including more SBEM datasets from more subjects and/or locations in the corpus callosum, as well as bigger sample size are necessary to increase our understanding of the 3D ultrastructure of the corpus callosum.

## Materials and Methods

### Animal model, tissue preparation, and SBEM imaging

#### Animals

Five adult male Sprague-Dawley rats (10-weeks old, weight 320 and 380g, Harlan Netherlands B.V., Horst, Netherlands) were used in the study. The animals were singly housed in a room (22 ±1 °C, 50% – 60% humidity) with 12 h light/dark cycle and free access to food and water. All animal procedures were approved by the Animal Care and Use Committee of the Provincial Government of Southern Finland and performed according to the guidelines set by the European Community Council Directive 86/609/EEC.

#### Traumatic brain injury model

TBI was induced by lateral fluid percussion injury in three rats (TBI-1, TBI-2, TBI-3)^62^. Rats were anesthetized with a single intraperitoneal injection (6mL/kg) of a mixture of sodium pentobarbital (58 mg/kg), magnesium sulphate (127.2mg/kg), propylene glycol (42.8%), and absolute ethanol (11.6%). A craniectomy (5mm in diameter) was performed between bregma and lambda on the left convexity (anterior edge 2.0 mm posterior to bregma; lateral edge adjacent to the left lateral ridge). Lateral fluid percussion injury was induced in one rat by a transient fluid pulse impact (21 ms) against the exposed intact dura using a fluid-percussion device (Amscien Instruments, Richmond, VA, USA). The impact pressure was adjusted to 3.1 atm to induce a severe injury. Two sham-operated rats (Sham-1, Sham-2) underwent all the same surgical procedures except for the impact.

#### Tissue processing

Five months after TBI or sham operation, the rats were transcardially perfused using 0.9% NaCl (30 mL/ min) for 2min followed by 4% paraformaldehyde (30 mL/ min) at 4 °C for 25min. The brains were removed from the skull and post-fixed in 4% paraformaldehyde /1% glutaraldehyde overnight at 4 °C.

#### Tissue preparation for SBEM

The brains were sectioned into 1-mm thick coronal sections with a vibrating blade microtome (VT1000s, Leica Instruments, Germany). From each brain, a section at approximately 3.80mm from bregma was selected and two samples from the ipsilateral and the contralateral corpus callosum were further dissected, as shown in Supplementary Fig. S6a. The samples were stained using an enhanced staining protocol^63^(see Supplementary Fig. S6b). First, the samples were immersed in 2% paraformaldehyde in 0.15 M cacodylate buffer containing 2mM calcium chloride (pH = 7.4), and then washed five times for 3 min in cold 0.15 M cacodylate buffer containing 2mM calcium chloride (pH = 7.4). After washing, the samples were incubated for 1 h on ice in a solution containing 3% potassium ferrocyanide in 0.3 M cacodylate buffer with 4 mM calcium chloride combined with an equal volume of 4% aqueous osmium tetroxide. They were then washed in double distilled H_2_O (ddH_2_O) at room temperature (5 × 3 min). Thereafter, the samples were placed in a solution of 0.01 mg/mL thiocarbohydrazide solution at room temperature for 20 min. The samples were then rinsed again in ddH2O (5 × 3 min), and placed in 2% osmium tetroxide in ddH2O at room temperature. Following the second exposure to osmium, the samples were washed in ddH_2_O (5 × 3 min), and then incubated in 1% uranyl acetate overnight at 4 °C. The following day, the samples were washed in ddH_2_O (5 × 3 min) and en *bloc* Walton’s lead aspartate staining was performed. In this step, the samples were incubated in 0.0066 mg/mL lead nitrate in 0.03 M aspartic acid (pH = 5.5) at 60 °C for 30 min, after which the samples were washed in ddH_2_O at room temperature (5 × 3 min), and dehydrated using ice-cold solutions of freshly prepared 20%, 50%, 70%, 90%, 100%, and 100% (anhydrous) ethanol for 5 min each, and finally placed in ice-cold anhydrous acetone at room temperature for 10 min. Embedding was performed in Durcupan ACM resin (Electron Microscopy Sciences, Hatfield, PA, USA). First, the samples were placed into 25% Durcupan^#1^ (without component C):acetone, then into 50% Durcupan^#1^:acetone, and after into 75% Durcupan^#1^:acetone overnight. The following day, they were placed in 100% Durcupan^#1^ for 2h in a 50 °C oven (2 times), and into 100% Durcupan^#2^ (4-component mixture) for 2 h in a 50 °C oven. Finally, the samples were embedded in 100% Durcupan^#2^ in Beem embedding capsules (Electron Microscopy Sciences) and baked in a 60 °C oven for 48 h.

After selecting the area within the samples, as shown in Supplementary Fig. S6c, the blocks were further trimmed into a pyramidal shape with a 1 × 1 mm^2^ base and an approximately 600 × 600 μm^2^ top (face), which assured the stability of the block while being cut in the SBEM microscope. The tissue was exposed on all four sides, bottom, and top of the pyramid. The blocks were then mounted on aluminum specimen pins using conductive silver epoxy (CircuitWorks CW2400). Silver paint (Ted Pella, Redding, CA, USA) was used to electrically ground the exposed block edges to the aluminum pins, except for the block face or the edges of the embedded tissue. The entire surface of the specimen was then sputtered with a thin layer of platinum coating to improve conductivity and reduce charging during the sectioning process.

#### SBEM data acquisition

All SBEM data were acquired on an SEM microscope (Quanta 250 Field Emission Gun; FEI Co., Hillsboro, OR, USA), equipped with the 3View system (Gatan Inc., Pleasanton, CA, USA) using a backscattered electron detector (Gatan Inc.). The top of the mounted block or face was the x-y plane, and the z direction was the direction of the cutting.

All the samples were imaged with a beam voltage of 2.5 kV and a pressure of 0.15 Torr. The datasets were acquired with a resolution of 13-18.3nm x 13-18.3nm x 50nm amounting to an area of 13.3-18.7μm x 13.3-18.7μm x 14.25μm in the x, y, and z directions, respectively. After imaging, Microscopy Image Browser^14^ was used to apply lateral registration to the slices. We quantified the registration using cross correlation analysis between successive slices and subtracting the running average (window size = 25) to preserve the directionality of axons while registration. Supplementary Fig. S6d shows a representative SBEM volume of the contralateral corpus callosum of the sham-operated rat. We also show two representative images cropped from the sham-operated and TBI volumes in Supplementary Fig. S6e and f, respectively.

### ACSON segmentation pipeline

The ACSON segmentation pipeline annotates the ultrastructure in SBEM volumes of white matter. The pipeline began with denoising of the SBEM volumes, and proceeded by segmenting the volumes using BVG. The segmented volumes were refined using supervoxel techniques, and, finally, the subcellular structures, cells, and myelinated and unmyelinated axons were annotated.

#### Denoising

SBEM images are degraded by noise from different sources, such as noise in the primary beam, secondary emission noise, and noise in the final detection system^64^. To suppress these complex noise patterns, we applied a non-local BM4D algorithm^65^. Unlike local averaging filters, which smooth an image by averaging values in the neighborhood of a target voxel, non-local filtering considers all the voxels in the image, weighted by how similar these voxels are to the target voxel. BM4D, in particular, enhances a sparse representation in the transform-domain by grouping similar 3D image patches (i.e., continuous 3D blocks of voxels) into 4D data arrays called, groups. The steps to realize the filtering are the 4D transformation of 4D groups, shrinkage of the transform spectrum, and inverse 4D transformation. While BM4D has been extensively used for denoising datasets from diverse imaging modalities, its application in 3D-SBEM datasets is novel. We applied the default parameter values of BM4D for denoising, which automatically estimates the noise type and variance. Figure 1a shows a slice of SBEM volume before filtering, and Fig. 1b illustrates the BM4D output in which the noise has been strongly attenuated. Segmentation. For the segmentation of a 3D-SBEM image, we devised a hybrid technique, named BVG, that integrates seeded region growing and edge detection. To elaborate BVG, we denote a 3D-SBEM image after denoising with BM4D as *z*(*x*): *X* → [0,1], where *x* ∈ *X* is a 3D spatial coordinate. Note that the intensity can range from 0 to 1. We defined *Ê* ⊂ *X* as the set of edges of *z*, using a Canny edge detector^66^. We set the parameter values of the Canny edge detector as follows: the SD of the Gaussian filter was 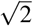 and thresholds for weak and strong edges were 0.25 and 0.6 times the maximum gradient magnitude. In SBEM volumes with resolution anisotropy and a coarser resolution in the *z* direction, regions in successive slices did not appear continuously, and areas close to the structure boundaries overlapped. Therefore, we dilated the set of edge coordinates *Ê* in-plane with a 3×3 square structuring element. The dilated edges are denoted as *E*. The edge dilation was proportional to the resolution anisotropy. We then used BVG to segment *z* into *n* + 1 distinct volumes *V*_1_, *V*_2_,…,*V_n_*, E, ⊂ *X*, in which 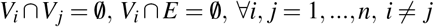. BVG is a serial segmentation algorithm, meaning that segmentation of *V_i_* starts only when *V*_-1_ is segmented. To segment *V_i_*, BVG begins with one voxel called the seed, denoted as *S_k_* ⊂ *V_i_*, which iteratively grows—*k* is the iteration number—and finally results in the volume *V_i_*. *N*(*S_k_*) is in the neighborhood of *S_k_* defined as 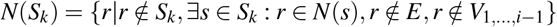, where *N*(*s*) is the 3D-neighborhood of voxel *s*. In each iteration, BVG appends a set of voxels *A* to *S_k_*, where *A* = {*x*|*x* ∈ *N*(*S_k_*), *δ*(*x*) ≤ *δ_T_*} and *δ*(*x*) measures the similarity of voxel *x* to the set *S_k_*. We defined the measure of similarity as 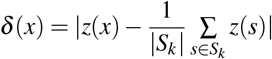 and set the similarity threshold *δ_T_* to 0.1. An iteration terminates, if 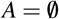, or |*S_k_*| ≥ *ϑ*, where *ϑ* is a volume constraint. If *V_i_* grew larger than *ϑ*, the results were discarded and the voxels within *Vi* were freed for other regions to compete for them.

The segmentation was initiated by annotating the low-intensity structures, i.e., myelin and mitochondria, which were considered together as *V*_1_. BVG was initiated with a random low-intensity voxel *S*_1_ with *z*(*S*_1_) ≤ 0.4. This one seed was sufficient to segment *V*_1_ because myelin is a connected structure in a consecutive SBEM image. We defined *N*(*s*) using 26 neighbors, and set *ϑ* = ∞. Figure 1c shows a slice of canny edges together with the segmented myelin and mitochondria (*V*_1_). To segment other structures *V*_2_, …, *V_n_*, we needed a more advanced seeding mechanism. Referring to Fig. 1c, we noticed that other structures are surrounded by myelin and edges. Therefore, we first generated a binary mask, *B*, of the union of the dilated edges *E* and the myelin-mitochondria segment *V*_1_ (Fig. 1c). Denoting each 2D-slice of *B* as *B_i_*, we computed the Euclidean distance transform^67^ for every *B_i_* individually, defined as *DT_i_* and shown in Fig. 1d. The pixel value in the distance transform *DT_i_* is the shortest distance from that pixel to a set of pixels *B_i_*. We defined the location of the seeds by extracting the regional maxima of each *DT_i_* (Fig. 1d). To segment *V_i_*, BVG was initiated with a seed from the set of extracted regional maxima not belonging to any previously segmented *V_j_, j* = 1,…, *i* – 1. We defined *N*(*s*) using 6 neighbors and set *ϑ* = 10^6^, which equals 12.5 μm^3^ of tissue or 1.5 times the volume of the largest axons in the dataset. Figure 1e shows one slice of the primary segmentation of the white matter ultrastructure, not belonging to *B*. The seed extraction overestimates the number of segmented volumes. This does not pose a problem, however, as the serial nature of BVG does not permit repetitive segmentation of an already segmented volume.

#### Segmentation post processing

The segmentation with BVG may result in small volumes, e.g., smaller than 5 × 10^3^ voxels, which actually belong to larger segments. As well, dilated Canny edges *E* should be assigned with a label as Fig. 1f shows. Therefore, we refined the segmentation by utilizing the SLIC supervoxel^68^ technique to relabel the small volumes and attach them to larger ones. Supervoxels group nearby voxels with similar intensity values into clusters. Particularly, SLIC clusters voxels based on a distance measure defined by 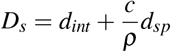, where *d_int_* ensures intensity similarity and *d_sp_* enforces voxel proximity to the supervoxels. In SLIC, the initial supervoxel centers are defined at regular grid steps *ρ*, and their compactness is controlled by *c*. Also, the spacing parameter *s* allows accounting for resolution anisotropy in *x, y*, and *z* directions. We assigned the SLIC arguments to produce compact and large supervoxels, while accounting for the resolution anisotropy. Thus, we set *c* = 23, *ρ* = 11 and *s* = [1,1, *vx*/*vz*], where *vx* and *vz* are the voxel size in *x* and *z* directions, respectively. Note that, voxel size in *x* and *y* directions was equal. Supplementary Fig. S7 shows the effect of altering *ρ, c* and *s* while generating supervoxels. We refined the large volumes *V_i_* with more than 5 × 10^3^ voxels by the SLIC supervoxels. In more detail, suppose that we have generated *Q* supervoxels *SV_q_, q* = 1,…, *Q*. Then, we re-defined *V_i_* as 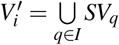 where 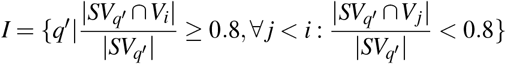. Refining the segmentation with SLIC technique eliminated most of the small volumes by relabeling and attaching them to the larger segments. Note that as the edges were included in the supervoxels, the supervoxel-based refinement also labeled voxels belonging to the edges (Fig. 1f).

#### Annotating subcellular structures and cells

The segmentation of myelin sometimes included mitochondria because the boundaries between these two structures were not clearly resolved. We wanted to label mitochondria separately, however, and include them as part of the myelinated axons. Not including mitochondria in the myelinated axon domain produces cavities, as shown in Fig. 1e,f. The cavities can be used to label the mitochondria. To detect the cavities in the myelinated axons, on each large volume, 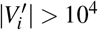, using a 3D distance transform, we propagated the surface of the volume for 1 μm. The surface of the enlarged volume was then propagated for – 1 μm shrinking of the volume. Applying this procedure to each large volume altered the morphology of the volume, and closed those cavities smaller than 1 μm. The difference between the altered volume and *V*′ was considered a potential mitochondrion, *M_i_*. We refined *M_i_* with SLIC supervoxels with the same parameter values and techniques mentioned in the segmentation post-processing section. Note that because some of the cavities were due to myelin, annotating the mitochondria was finalized using human supervision to check for myelin. Figure 1g shows the final result of the mitochondria segmentation. The myelin segment was then re-defined as the set difference of *V*_1_ and all mitochondria, denoted as *MY*. In our SBEM-datasets, vacuoles appeared brighter than all of the other ultrastructures. Thus, if 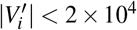 and 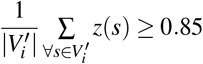, we labeled 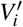 as a vacuole. We defined the remaining volumes, not mitochondria nor vacuole, as axons denoted as *AX_i_, i* = 1,…, *m*. To distinguish if an axon *AX_i_* was a myelinated or an unmyelinated axon, we studied a thick hollow cylinder enclosing the axon. The enclosing cylinder was formed by those supervoxels having a common face with the axon *AX_i_*. If the enclosing cylinder contained myelin above a threshold, the axon has been considered as a myelinated axon. In more detail, let *Λ_i_*, be the indexes of supervoxels enclosing an axon *AX_i_*. If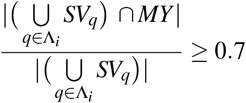, we considered *AX_i_* to be a myelinated axon. Note that because unmyelinated axons can be surrounded by several myelinated axons, they can be miss-classified as myelinated axons, thus requiring a proof reading after the classification.

To label cells and cell-processes, we considered a straightforward approach as the volumes of cells were expected to be larger than the volumes of any other structure, excluding myelin. Recall that we set the volume threshold *ϑ* = 10^6^ for the segmentation of *V*_2_,…, *V_n_*, which leaves some voxels unlabeled. These unlabeled voxels *X*′ comprised cells and cell processes. We segmented *X*′ into *n*′ cells using connected component analysis. In general, we detected 1-4 cell bodies/process in each SBEM-volume. Figure 1h demonstrates the final segmentation results of myelin, myelinated axons, unmyelinated axons, oligodendrocyte cell body and its processes. Mitochondria and vacuoles belonging to myelinated axons were colored the same as their corresponding myelinated axons. Figure 1i and Supplementary Fig. S4 show the 3D rendering of myelinated axons in contralateral corpus callosum of Sham-1 dataset.

### ACSON morphometry pipeline

We defined a cross-section as the intersection of a segmented myelinated axon Ω and a perpendicular plane to the axonal skeleton *γ*^69^. To detect the myelinated axon skeleton *γ* with sub-voxel precision, we adapted a method from Van Uitert & Bitter^70^. First, we defined three points in the myelinated axon domain Ω: *x*\* with the largest distance from the myelinated axon surface Γ, and *x*_*e*1_ and *x*_*e*2_ as the endpoints of the axons, i.e., the tips of the axon. The minimum-cost path connecting *x*_*e*1_ to *x*_*e*2_ through *x*\* was the axon skeleton γ. The path was found in two steps, first from *x*_*e*1_ to *x*\*, and then from *x*_*e*2_ to *x*\*. Mathematically,

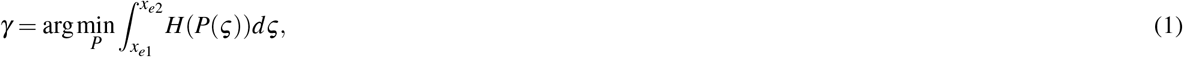

where *ς* traces the path *P*, and *H* is the cost function. To enforce the minimum-cost path to run at the middle of the object, the cost function *H* should be higher if the path moves away from the center. Points *x*\*, *x*_*e*1_, and *x*_*e*2_ and solving equation (1) was defined by solving an eikonal equation on the axonal domain Ω. The eikonal equation is a non-linear partial differential equation defined as a special case of wave propagation in which the front Γ advances monotonically with speed *F*(*x*) > 0. The eikonal equation can be formulated as |∇*T*(*x*)|*F*(*x*) = 1, where *T*|_Γ_ = 0. The solution, *T*(*x*), is the shortest time needed to travel from Γ to any point *x* ∈ Ω, with the speed *F*(*x*) > 0. Although the eikonal equation can be solved with the FMM^71^, we used 3D MSFM^51^. MSFM combines multiple stencils and second-order approximation of the directional derivatives over the FMM to improve the accuracy of solving the eikonal equation on Cartesian domains. To find *x*\*, we computed the time-crossing map *T*_1_(*x*) from the myelinated axon interface Γ with constant speed *F*_1_(*x*) = 1, *x* ∈ Ω. The global maximum of *T*_1_, where *T*_1_ (*x*) ≤ *T*_1_ (*x*\*), *∀x* ∈ Ω was defined as *x*\*. To find *x*_*e*1_, *x*_*e*2_, and *γ*, we calculated a new time-crossing map *T*_2_ (*x*), starting at *x*\* to every voxel in Ω, with a non-constant speed 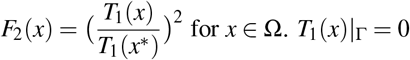. Using *H*(*x*) = 1 – *F*_2_(*x*) to define the cost ensured that voxels in the middle of the myelinated axon were reached prior to the voxels close to Γ. We defined the furthest point from *x*\* on the *T*_2_ map, i.e., the global maximum of *T*_2_, as *x*_*e*1_. Similarly, *x*_*e*2_ was defined as the furthest point from *x*_*e*1_, at the global maximum of the time-crossing map *T*_3_(*x*), starting from *x*_*e*1_ to every voxel in Ω, with speed *F*_2_(*x*). For both endpoints, we determined the minimum-cost path between *x*_*ei*_ and *x*\*, *γ_i_, i* = 1,2, by backtracking, starting from *x_ei_* and progressing along the negative gradient 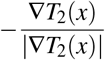 until *x*\* was reached. *x*\* is the global minimum of *T*_2_(*x*), so that we were guaranteed to find it with backtracking. The backtracking procedure can be described by the ordinary differential equation 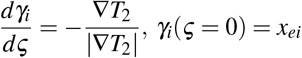, where *ς* traces *γ_i_*. We used a 4^th^ order Runge-Kutta scheme, with a 25 nm step size, to solve the ordinary differential equation with sub-voxel accuracy. The myelinated axon skeleton was formed as *γ* = *γ*_1_ ∪ *γ*_2_. Note that computing the skeleton in this way prevented the skeleton from cutting corners^72^. Figure 3a shows a 3D reconstruction of a myelinated axon, its mitochondria, and the extracted skeleton (axonal axis). Note that *x*_*e*1_ and *x*_*e*2_ defined as the global maxima of *T*_2_(*x*) and *T*_3_(*x*), lie on the myelinated axon surface Γ, and not in the center of the myelinated axon. The cost function H, however, forces the skeleton to immediately move away from the surface Γ toward the center. Therefore, we dropped the first 1 μm at both ends of *γ* in our later calculations.

To determine the cross-sectional planes perpendicular to *γ*, we formed a moving reference frame of the size 8 μmx8 μm with 50 nm resolution. At each skeleton point *ς*, the unit tangent vector to *γ* was used to define the orientation of the reference frame. The intersection of the reference frame with the myelinated axon defined the cross-section of the myelinated axon.

The intensity values of a cross-section ranged between 0 and 1. Each cross-section was thresholded at 0.5, resulting in a 2D binary image *C* denoted as *C*: *X* → {0,1}, where the point *x* = (*x*_1_,*x*_2_) was foreground iff *x* ∈ *C*. We defined the center point of *C* as 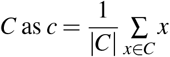. By translating the binary 2D cross-section *C* to the center of 2D Cartesian coordinate *C_t_* = {*y* ∈ *X* : *y* = *x* – *c*}, we found an ellipse that had the same normalized second central moment as *C_t_*. The cross-sectional morphology of myelinated axons were quantified by computing the minor and major axes and the eccentricity of the fitted ellipse^73^, and the diameter of a circle with the same area as the cross-section, called equivalent diameter.

### Evaluation of segmentation accuracy

#### Manual segmentation

The manual segmentation by A.S. defined each ultrastructure as its own region, i.e., different axons had distinct labels in the manual segmentation as in the automatic one. It also annotated each segmented region as myelin, myelinated or unmyelinated axon. In the annotated images, mitochondria and vacuoles were included into intra-axonal space.

#### Precision and recall

For a tissue-type level evaluation, we used the precision and recall as in the previous studies^28,32^. Let A and B be the sets of voxels of a particular tissue-type (myelin, myelinated axon, unmyelinated axon) in the manual and automated segmentations, respectively. We defined 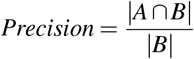, and 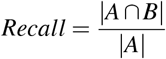. The maximum for the precision and recall is equal to one when the automated segmentation perfectly matches the manual segmentation. These metrics do not describe topological differences between the manual and automated segmentations. For example, these metrics do not penalize the automatic segmentation for incorrectly dividing a single axon into two axons.

#### Weighted Jaccard index and weighted Dice coefficient

To further evaluate the automated segmentation, we used Jaccard index and Dice coefficients in the region level. The Jaccard index^74^ and Dice coefficient^75^ is defined by 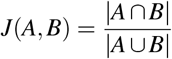 and *Dice Coef* 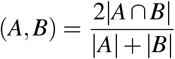, where *A* and *B* are the regions segmented manually and automatically, respectively. The maximum for these metrics is equal to one occurring when *A* perfectly matches *B*. If no overlap occurs between *A* and *B*, these metrics are equal to zero. Let *A_i_, i* = 1,…, *a*′, and *B_j_, j* = 1,…, *b*′ be the regions in the manual and automated segmentation, respectively. To assign *A_i_*, and the best matching *B_j_*, we formed a similarity matrix based on Dice coefficients for any possible pair of *A_i_*, and *B_j_*, where the element *(i, j)* of the similarity matrix was *Dice Coef* (*A_i_*;, *B_j_*). We used the Hungarian algorithm^76,77^ to match the regions. We defined the weighted mean of the Jaccard index and the weighted mean of the Dice coefficients as 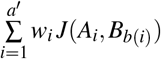 and 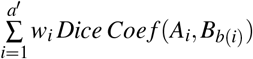, respectively, where *b*(*i*) is the index of the region best matching *A_i_* and the weight 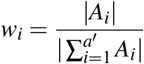.

### Comparison of 2D and 3D morphological analyses

To simulate 2D morphometry, we assumed that the myelinated axon morphometry is quantified on a single image of the 3D image stack. For each myelinated axon, we determined the best orientation of the image stack for the 2D quantifications by extracting the Euler angles of a fitted ellipsoid to the segmented myelinated axon. We randomly selected a single image in that orientation to present the myelinated axon. For example, if a myelinated axon was elongated parallel to the *z*-axis, we selected a random image parallel to the *x-y* plane. Each myelinated axon was quantified separately for its minor and major axes, equivalent diameter, and eccentricity. The relative difference between the 2D and 3D quantifications for each myelinated axon was defined as 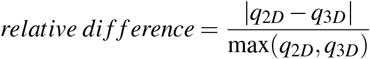, where *q* is the quantity of interest, i.e., minor and major axes, equivalent diameter, and the eccentricity, measured by the 2D or 3D procedures. Note that, for the 3D morphometry, median of the measurements along the axonal axis were quantified Fig. 4a.

### Statistical analysis

#### Nested ANOVA

Nested (hierarchical) ANOVA is a parametric hypothesis testing and an extension of 1-way ANOVA. A nested ANOVA is used when there is one measurement variable and more than one nominal variable, and the nominal variables are nested^35^. The nominal variables being nested means that each value of one nominal variable (the subgroups) is found in combination with only one value of the higher-level nominal variable (the groups). We considered lower-level variables (cross-section, axon, animal) as random effects variables and the top level (group, either sham or TBI) as a fixed effect variable. The null hypotheses were whether there existed significant variation in means among groups at each level. The analysis was performed using the anovan function of MATLAB R2017b, with type II sum of squares. Because, the design matrix of anovan grows quickly when the nesting levels increases, we measured the median of the cross-sectional quantities and assigned them to the lowest level of nested ANOVA. At the cross-sectional level, the distributions of equivalent diameter, minor and major axes and eccentricity were multimodal (Supplementary Fig. S5), thus median of cross-sections was preferred to mean. The analysis was performed separately for two hemispheres.

#### Variance components

Nested ANOVA partitions the variability of the measurements into different levels. The variance components describe what percentage of the total variance is attributable to each level.^35^.

## Acknowledgements

This work was supported by the Academy of Finland (J.T. and A.S.), and Biocenter Finland and University of Helsinki (I.B., E.J., and SBEM imaging). We would like to thank Maarit Pulkkinen for her help with the animal handling, and Mervi Lindman and Antti Salminen for help preparing the samples for SBEM.

## Author contributions

A.A., J.T., and A.S. conceived the project and designed the study. A.A. implemented the methods and performed the experiments. A.A., J.T., and A.S. analyzed the data. I.B. and E.J. contributed to electron microscopy imaging. A.A., J.T., and A.S. wrote the manuscript. All authors commented on and approved the final manuscript.

## Availability of data and material

The datasets used and/or analyzed during the current study are available from the corresponding author on reasonable request. The source code of ACSON is available at https://github.com/aAbdz/ACSON.

## Supplementary Information

**Figure S1.**
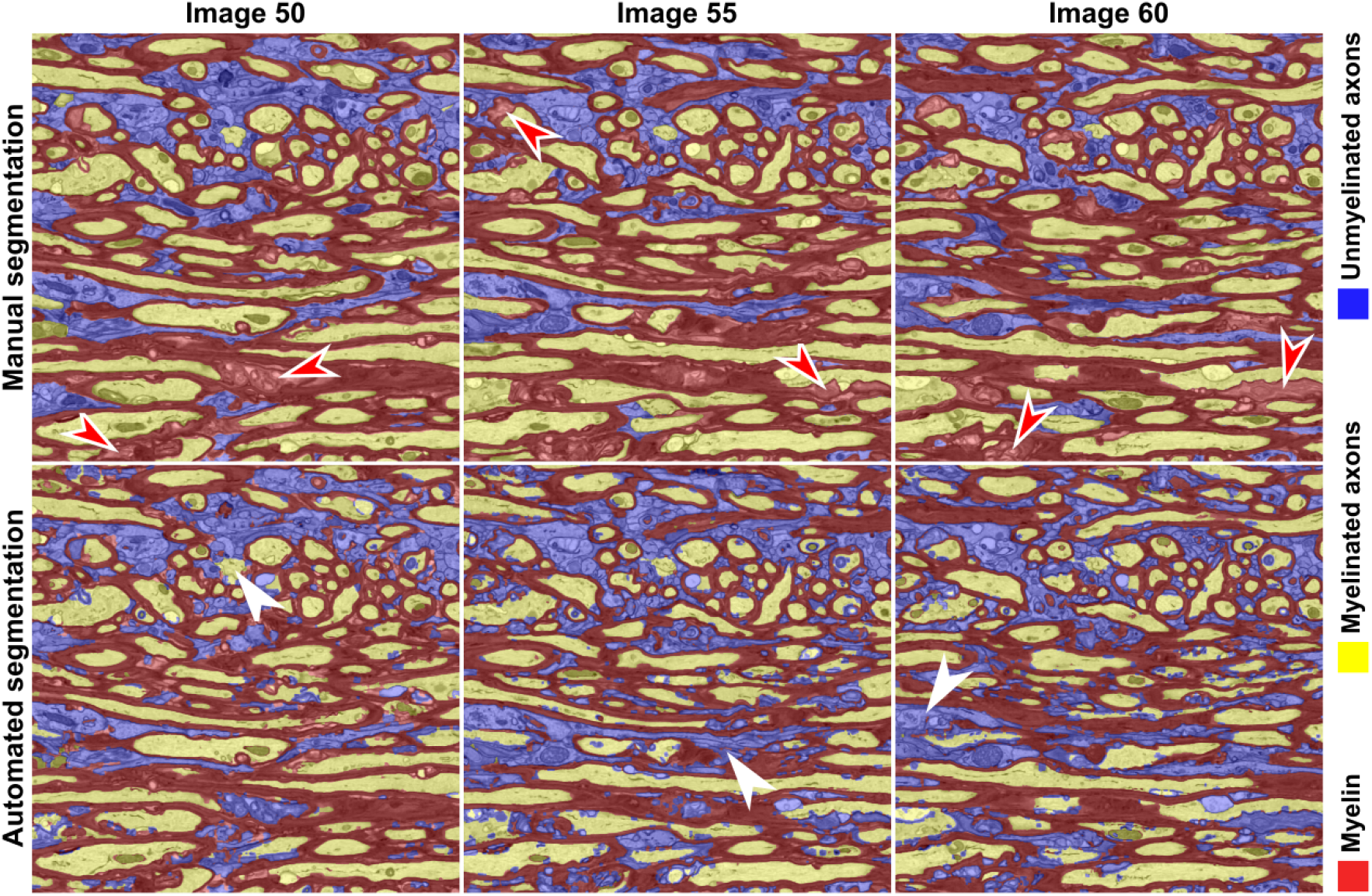
Manual and automated segmentations of three images from the contralateral corpus callosum of Sham-1 dataset, represented with a 3 labels map of myelin and intra-axonal space of myelinated and unmyelinated axons. As in Fig. 2, the red arrowheads indicate delamination in the myelin sheaths. These substructures were annotated as myelin in the manual annotation, while the automated segmentation excluded the myelin delamination from the myelin-labeled structures. White arrowheads point at segmentation errors where the membrane of unmyelinated axons was poorly resolved.

**Figure S2.**
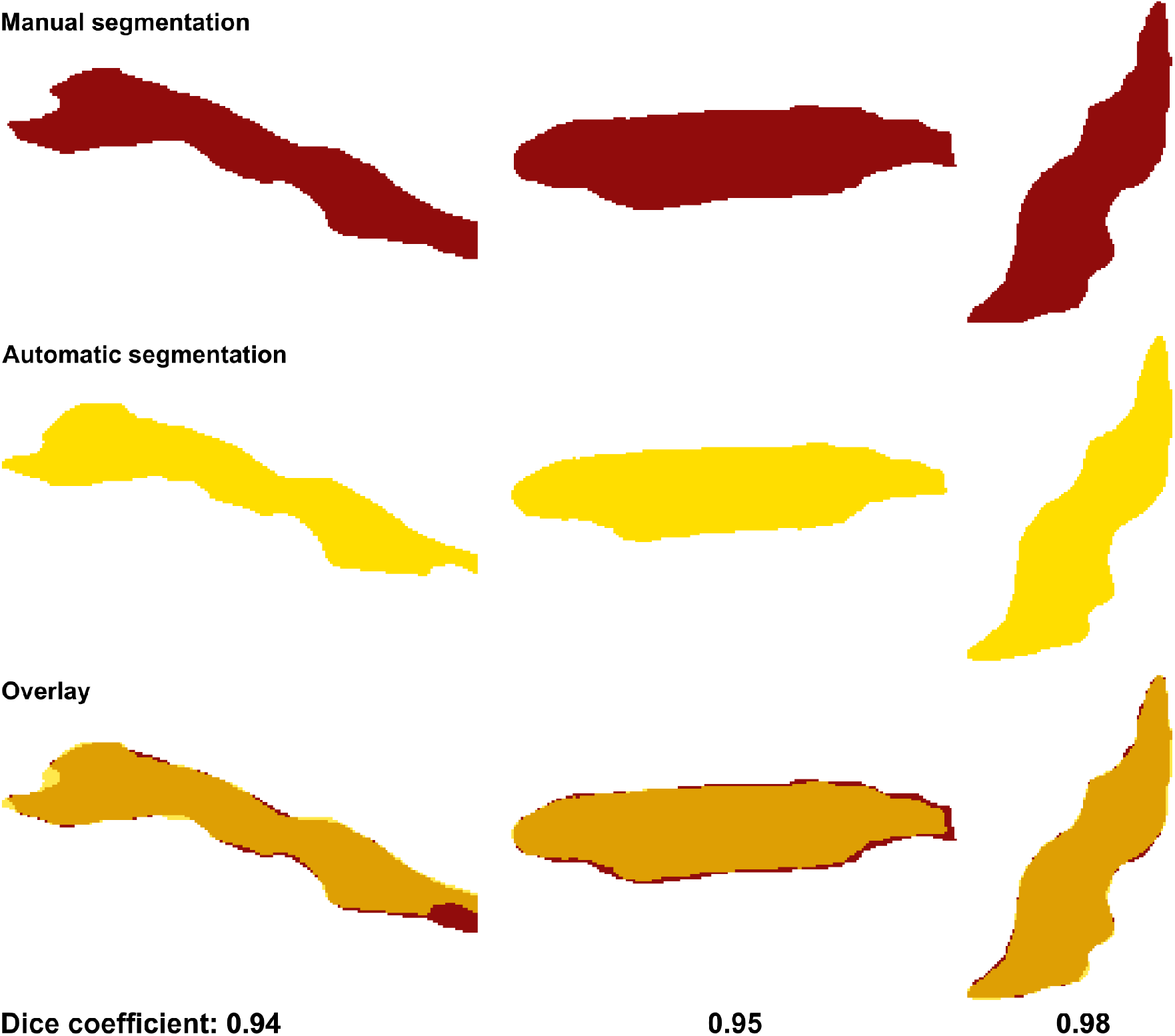
Comparison of manual and automated segmentation. Three representative segments of myelinated axons selected from contralateral corpus callosum of Sham-1 dataset. The Dice coefficients are sensitive to minor displacements in the location of boundaries.

**Figure S3.**
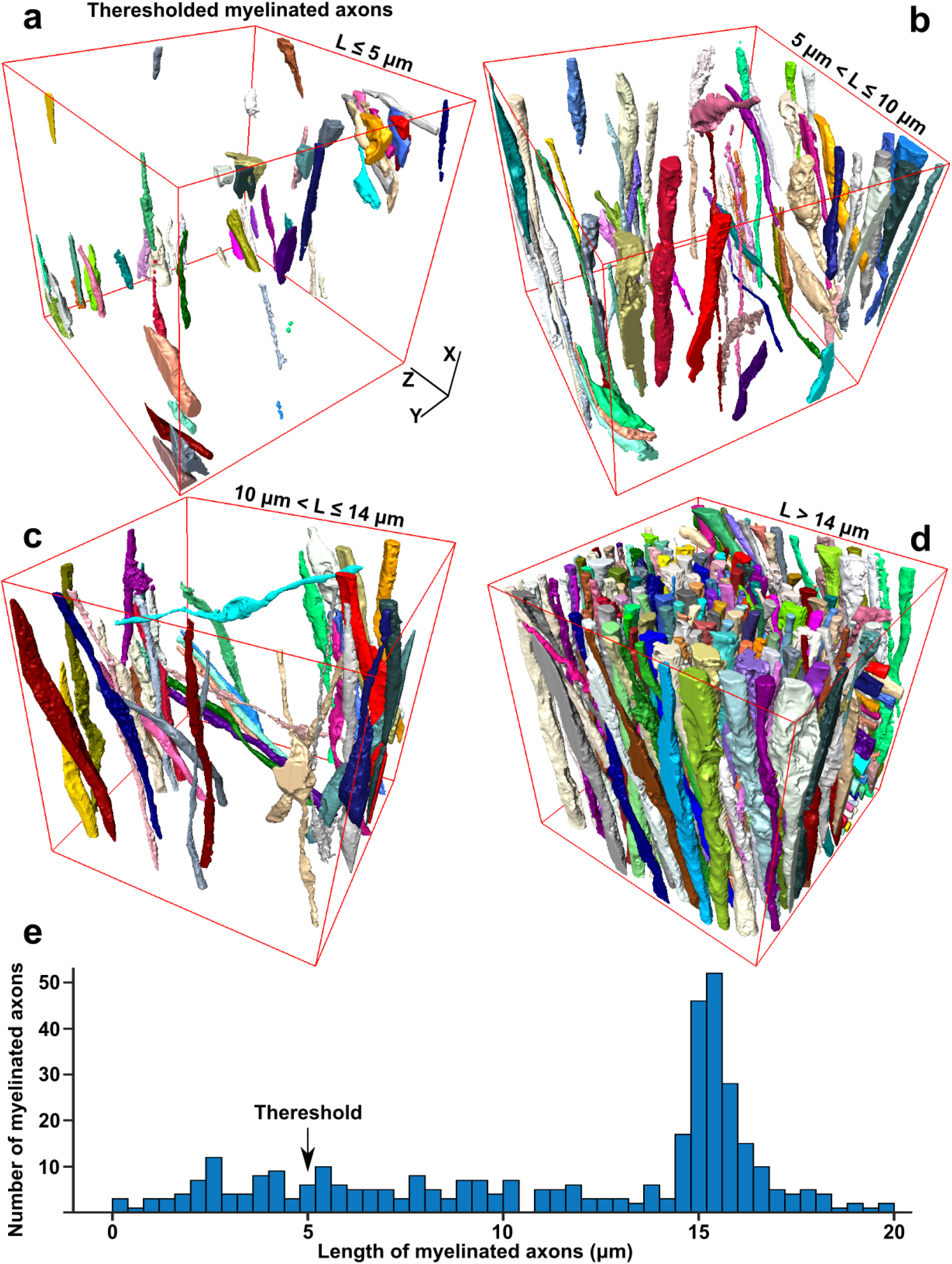
Evaluation of split and merge errors in segmentation of myelinated axons in contralateral corpus callosum of Sham-1 dataset. (a-d) 3D rendering of myelinated axons represented based on their length (L). Short axons (L ≥ 5 μm) traversed the corners of the SBEM volume, and they were not the result of split error. (**e**) Distribution of the length of myelinated axons. We thresholded myelinated axons at 5 μm to exclude myelinated axons traversing partially the SBEM volumes from later quantification. About 17% of myelinated were shorter than the threshold.

**Figure S4.**
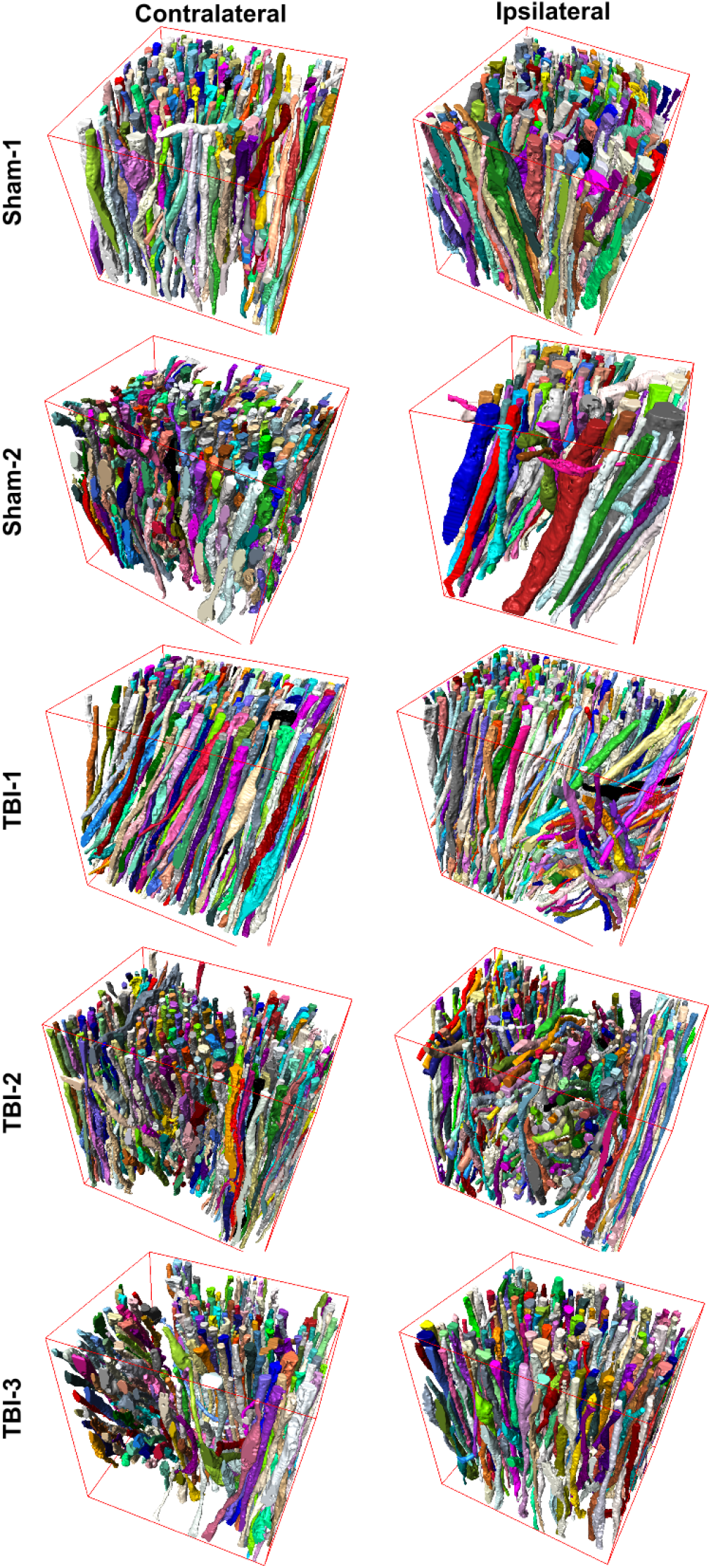
3D rendering of myelinated axons for all SBEM datasets. Myelinated axons with different thicknesses running along the volumes and organizing bundles with different orientations.

**Figure S5.**
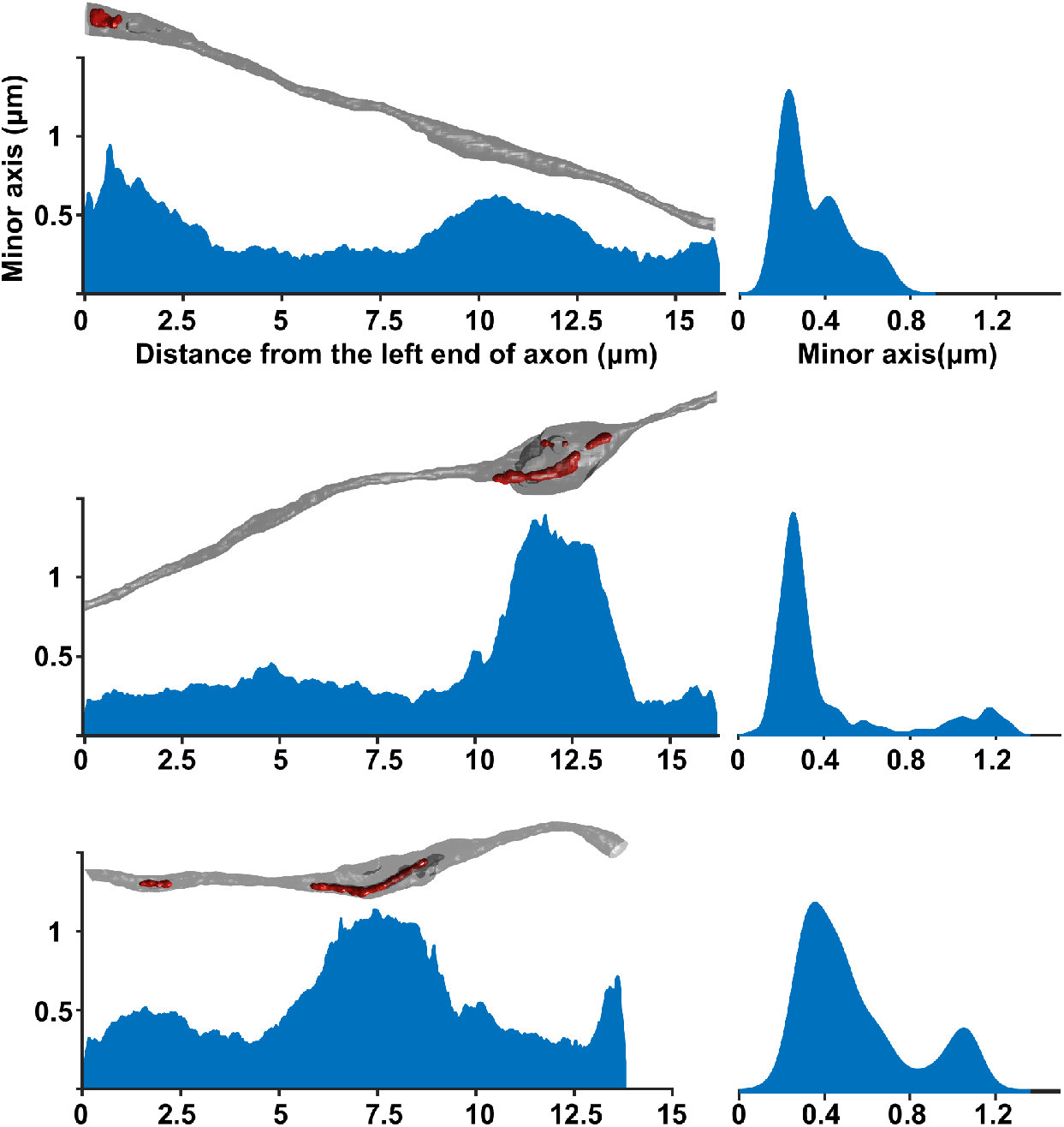
A cross-sectional diameter (minor and major axes or equivalent diameter) is not a constant parameter along a myelinated axon. Accumulation of organelles, such as mitochondria, increases the cross-sectional diameter locally. Therefore, the histogram of cross-sectional diameters for a myelinated axon is more likely to be bimodal.

**Figure S6.**
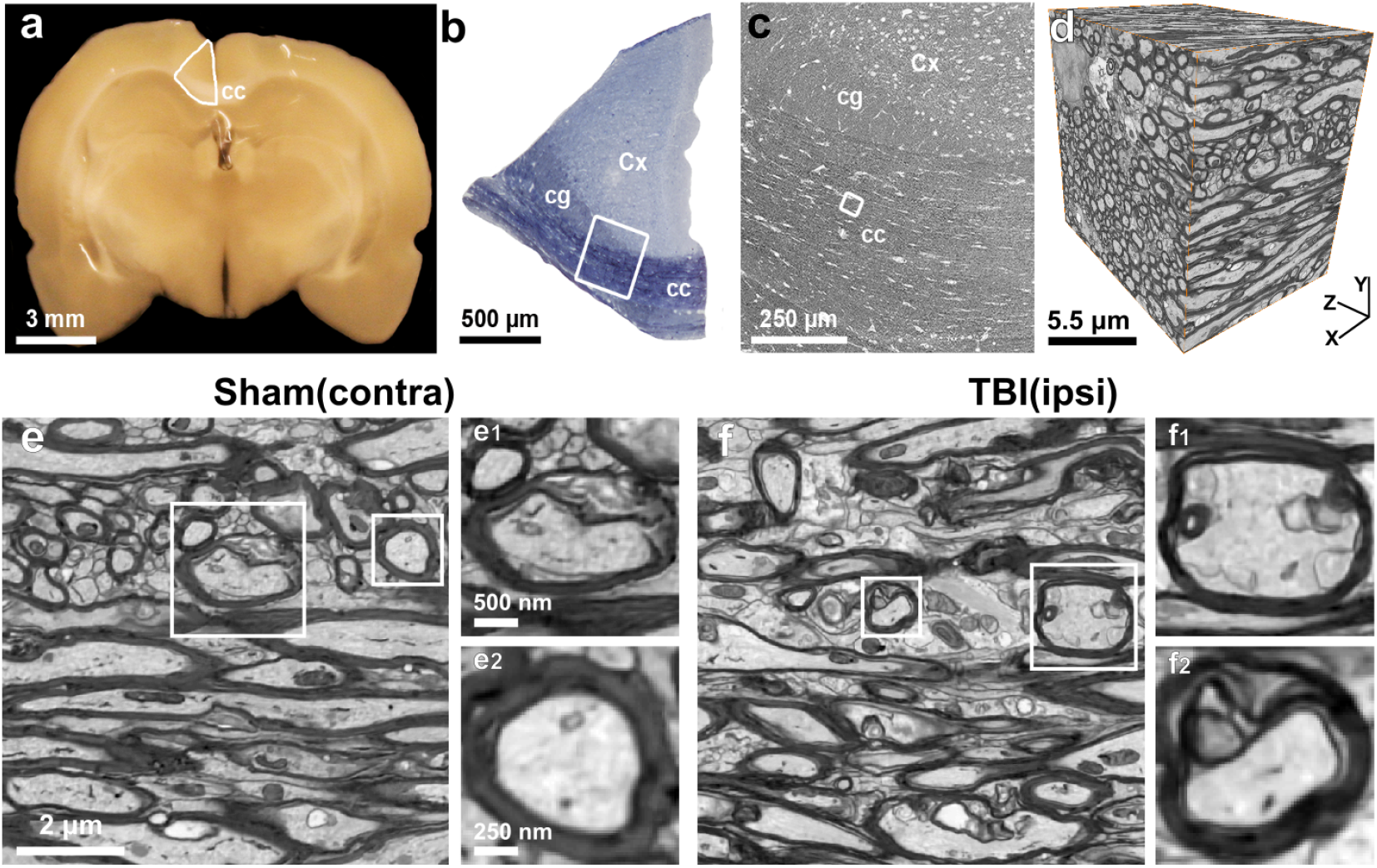
(**a**) A photomicrograph of 1 mm thick coronal section of a sham-operated (Sham-1) rat brain. The white outline shows the selected section for staining, containing part of the corpus callosum, cingulum, and the cerebral cortex. (**b**) A photomicrograph of a semithin section stained with toluidine blue. The white outline shows the block trimmed for SBEM imaging. (**c**) Selection for SBEM imaging (white outline). (**d**) The SBEM volume of the contralateral corpus callosum of Sham-1 dataset. (**e**) A representative 2D image from **d**. (**e1**) and (**e2**) are two representative myelinated axons cropped from **e**. (**e1**) shows myelin delamination. (**f**) A representative 2D image of the ipsilateral corpus callosum of TBI-3 dataset. (**f1**) and (**f2**) are two representative myelinated axons from **f** indicating more frequent myelin delamination as the result of the induced severe injury.

**Figure S7.**
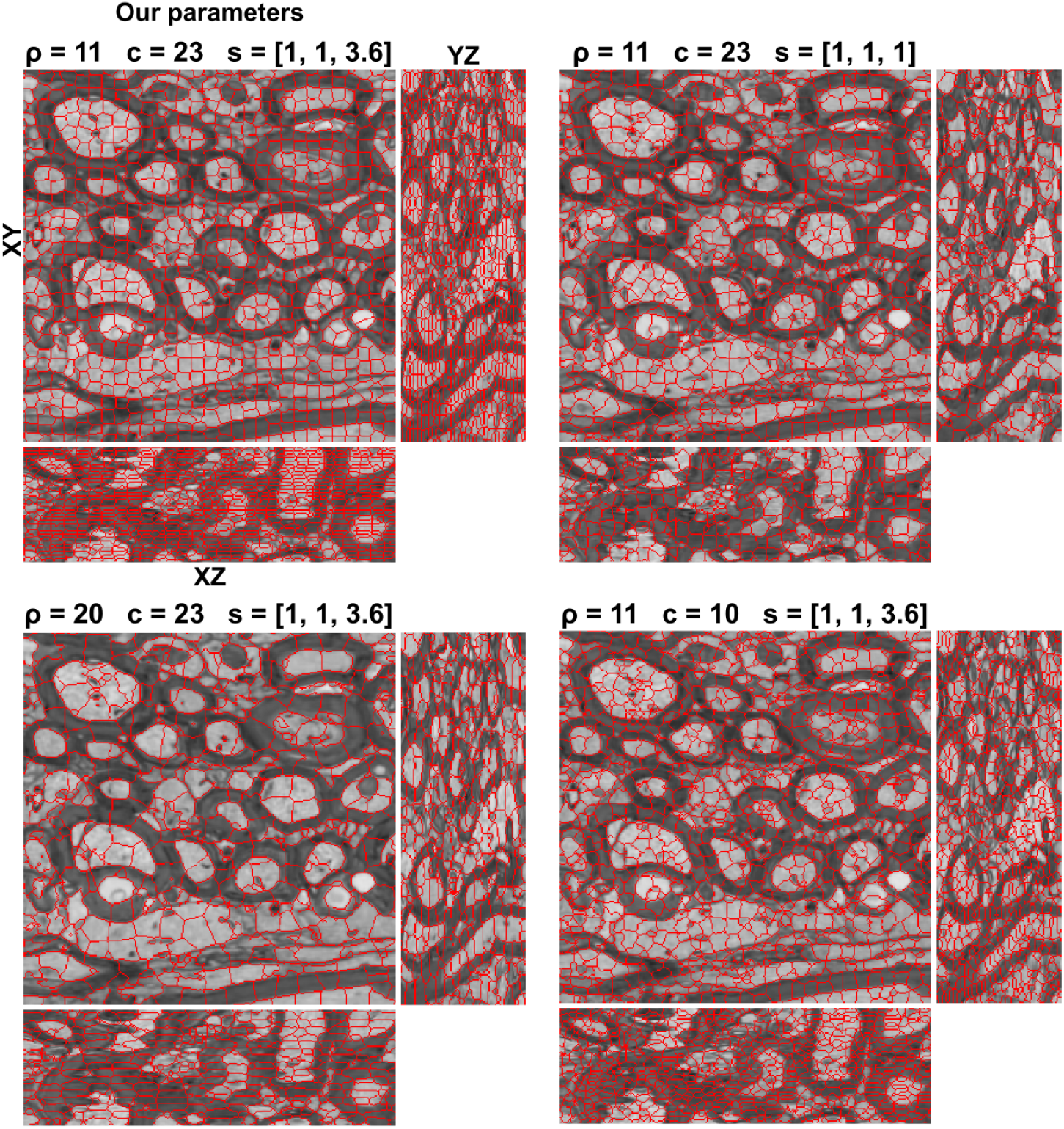
Supervoxel size, compactness and spacing as a function of parameters *ρ, c* and *s* for a cropped representative SBEM image. Spacing parameter *s* allowed us to account for the resolution anisotropy (coarser resolution in *z* direction) for generating supervoxels. We set spacing parameter in *z* direction 3.6 times bigger than *x* and *y* directions, as the voxel size of Sham-1 dataset was 13.8 nm x 13.8 nm x 50nm. Increasing *ρ* enlarges the supervoxels and decreasing *c* produces more compact and regular supervoxels.

**Table S1.**
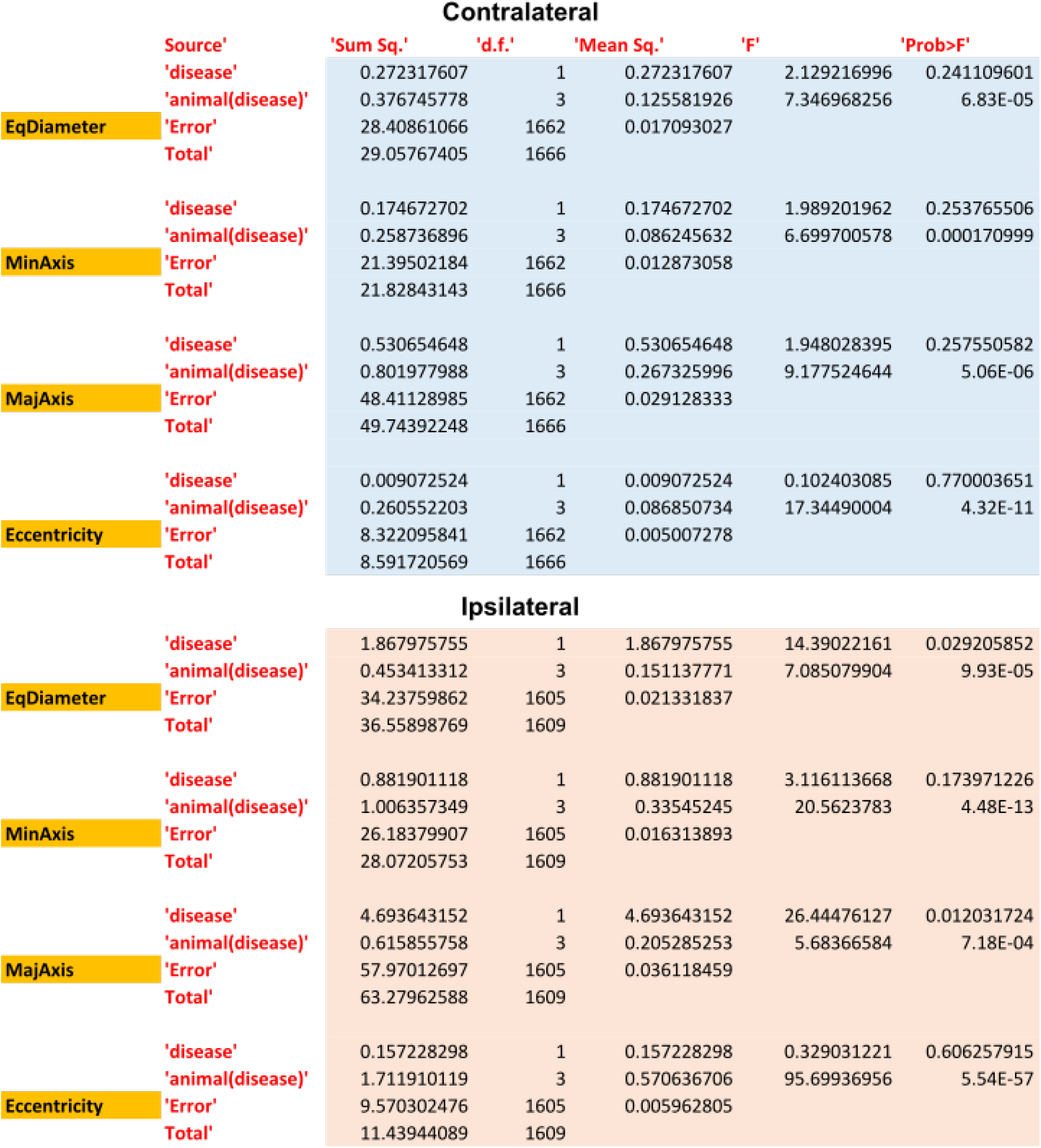
Nested ANOVA tables related to Fig. 5 of the equivalent diameter, the minor and major axes and the eccentricity in the contralateral and ipsilateral corpus callosum. Median of cross-sectional measurements was assigned to the lowest level nested ANOVA.

**Table S2.** Relative volume of the main cellular components to the SBEM volume, expressed as a percentage. Results are dataset-dependent, as the volume fraction that a cell body/process occupies varies among datasets. This affects the volume fraction of the ultrastructure in a dataset, preventing direct cross-analysis between datasets. Therefore, we calculated the aggregate g-ratio and the density of myelinated axons and mitochondria.

|  | Sham-1 |  | Sham-2 |  | TBI-1 |  | TBI-2 |  | TBI-3 |  |
| --- | --- | --- | --- | --- | --- | --- | --- | --- | --- | --- |
|  | Contra | Ipsi | Contra | Ipsi | Contra | Ipsi | Contra | Ipsi | Contra | Ipsi |
| Myelin (%) | 53.5 | 57.3 | 39.8 | 45.3 | 54.5 | 44.9 | 59.5 | 58 | 52 | 51.7 |
| Myelinated axons (%) | 26.9 | 31.4 | 34.4 | 25.5 | 26.9 | 23.1 | 23.8 | 25.4 | 27 | 30.6 |
| Number of myelinated axons | 367 | 404 | 548 | 234 | 459 | 541 | 650 | 851 | 534 | 418 |
| Mitochondria (%) | 3.1 | 4.2 | 2.2 | 1.6 | 3.7 | 4.9 | 3.1 | 3.5 | 2.4 | 2.6 |
| Vacuoles(%) | 0.9 | 1.6 | 1.4 | 0.1 | 0.5 | 0 | 1.7 | 0.3 | 1.5 | 1.2 |
| Unmyelinated axons (%) | 15.5 | 9.5 | 3.7 | 10.8 | 9.3 | 12.8 | 7.8 | 13.1 | 14.5 | 16.4 |
| Cells (%) | 4.1 | 1.8 | 22.1 | 18.4 | 9.3 | 19.2 | 8.9 | 3.5 | 6.5 | 1.3 |
| Aggregate g-ratio | 0.58 | 0.60 | 0.68 | 0.60 | 0.58 | 0.58 | 0.53 | 0.55 | 0.58 | 0.61 |
| Density of ultrastructure |  |  |  |  |  |  |  |  |  |  |
| Myelinated axons ($\mu\text{m}^{-3}$) | 0.13 | 0.12 | 0.14 | 0.11 | 0.15 | 0.13 | 0.14 | 0.17 | 0.14 | 0.10 |
| Mitochondria ($\mu\text{m}^{-3}$) | 0.63 | 0.58 | 0.65 | 1.12 | 0.22 | 0.79 | 0.12 | 0.16 | 0.28 | 0.51 |

## Supplementary video file

The video shows the segmentation of myelin, cell and myelinated axons in the contralateral corpus callosum of sham-1 dataset, respectively. Access/download the supplementary video at: http://www.doi.org/10.5281/zenodo.1459612.

## References

1. Guillery, R. W. Light- and electron-microscopical studies of normal and degenerating axons. In Contemporary Research Methods in Neuroanatomy, 77–105 (Springer Berlin Heidelberg, Berlin, Heidelberg, 1970). URL http://www.springerlink.com/index/10.1007/978-3-642-85986-1_5. DOI 10.1007/978-3-642-85986-1_5.

2. Wilkins, A. et al. Slowly progressive axonal degeneration in a rat model of chronic, nonimmune-mediated demyelination. J. Neuropathol. & Exp. Neurol. 69, 1256–1269 (2010). URL https://academic.oup.com/jnen/article-lookup/doi/10.1097/NEN.0b013e3181ffc317. DOI 10.1097/NEN.0b013e3181ffc317.

3. Lakovic, K. et al. Bilirubin and its oxidation products damage brain white matter. J. Cereb. Blood Flow & Metab. 34, 1837–1847 (2014). URL http://journals.sagepub.com/doi/10.1038/jcbfm.2014.154. DOI 10.1038/jcbfm.2014.154.

4. Donovan, V. et al. Repeated mild traumatic brain injury results in long-term white-matter disruption. J. cerebral blood flow metabolism 34, 715–723 (2014). URL http://www.ncbi.nlm.nih.gov/pubmed/24473478. DOI 10.1038/jcbfm.2014.6.

5. Torrealba, F. & Carrasco, M. A. A review on electron microscopy and neurotransmitter systems. Brain Res. Rev. 47, 5–17 (2004). DOI 10.1016/j.brainresrev.2004.06.004.

6. Denk, W. & Horstmann, H. Serial block-face scanning electron microscopy to reconstruct three-dimensional tissue nanostructure. PLoS Biol. 2 (2004). DOI 10.1371/journal.pbio.0020329.

7. Harris, K. M. et al. Uniform serial sectioning for transmission electron microscopy. J. Neurosci. 26, 12101–12103 (2006). URL http://www.jneurosci.org/cgi/doi/10.1523/JNEUROSCI.3994-06.2006. DOI 10.1523/JNEUROSCI.3994-06.2006.

8. Hayworth, K., Kasthuri, N., Schalek, R. & Lichtman, J. Automating the collection of ultrathin serial sections for large volume TEM reconstructions. Microsc. Microanal. 12, 86–87 (2006). URL http://www.journals.cambridge.org/abstract_S1431927606066268. DOI 10.1017/S1431927606066268.

9. Knott, G., Marchman, H., Wall, D. & Lich, B. Serial section scanning electron microscopy of adult brain tissue using focused ion beam milling. J. Neurosci. 28, 2959–2964 (2008). URL http://www.jneurosci.org/cgi/doi/10.1523/JNEUROSCI.3189-07.2008. DOI 10.1523/JNEUROSCI.3189-07.2008.

10. Helmstaedter, M., Briggman, K. L. & Denk, W. 3D structural imaging of the brain with photons and electrons. Curr. Opin. Neurobiol. 18, 633–641 (2008). DOI 10.1016/j.conb.2009.03.005.

11. Briggman, K. L. & Bock, D. D. Volume electron microscopy for neuronal circuit reconstruction. Curr. Opin. Neurobiol. 22, 154–161 (2012). URL http://dx.doi.org/10.1016/j.conb.2011.10.022. DOI 10.1016/j.conb.2011.10.022.

12. Helmstaedter, M., Briggman, K. L. & Denk, W. High-accuracy neurite reconstruction for high-throughput neuroanatomy. Nat. Neurosci. 14, 1081–1088 (2011). URL http://www.nature.com/doifinder/10.1038/nn.2868. DOI 10.1038/nn.2868.

13. Cardona, A. et al. TrakEM2 software for neural circuit reconstruction. PLoS ONE 7 (2012). DOI 10.1371/journal.pone.0038011.

14. Belevich, I., Joensuu, M., Kumar, D., Vihinen, H. & Jokitalo, E. Microscopy Image Browser: A platform for segmentation and analysis of multidimensional datasets. PLoS Biol. 14, 1–13 (2016). DOI 10.1371/journal.pbio.1002340.

15. Saalfeld, S., Cardona, A., Hartenstein, V. & Tomančák, P. CATMAID: Collaborative annotation toolkit for massive amounts of image data. Bioinforma. 25, 1984–1986 (2009). DOI 10.1093/bioinformatics/btp266.

16. Kaynig, V. et al. Large-scale automatic reconstruction of neuronal processes from electron microscopy images. Med. Image Analysis 22, 77–88 (2015). DOI 10.1016/j.media.2015.02.001.

17. Sommer, C., Straehle, C., Ullrich, K. & Hamprecht, F. a. ILASTIK : Interactive learning and segmentation toolkit. Eighth IEEE Int. Symp. on Biomed. Imaging (ISBI) 230–233 (2011). DOI 10.1109/ISBI.2011.5872394.

18. Berning, M., Boergens, K. M. & Helmstaedter, M. SegEM: Efficient image analysis for high-resolution connectomics. Neuron 87, 1193–1206 (2015). URL http://dx.doi.org/10.1016/j.neuron.2015.09.003. DOI 10.1016/j.neuron.2015.09.003.

19. Andres, B., Köthe, U., Helmstaedter, M., Denk, W. & Hamprecht, F. A. Segmentation of SBFSEM volume data of neural tissue by hierarchical classification. Lect. Notes Comput. Sci. (including subseries Lect. Notes Artif. Intell. Lect. Notes Bioinformatics) 5096 LNCS, 142–152 (2008). DOI 10.1007/978-3-540-69321-5_15.

20. Chklovskii, D. B., Vitaladevuni, S. & Scheffer, L. K. Semi-automated reconstruction of neural circuits using electron microscopy. Curr. Opin. Neurobiol. 20, 667–675 (2010). URL http://dx.doi.org/10.1016/j.conb.2010.08.002. DOI 10.1016/j.conb.2010.08.002.

21. Jain, V. et al. Learning to agglomerate superpixel hierarchies. Adv. Neural Inf. Process. Syst. 648–656 (2011). URL http://papers.nips.cc/paper/4249-learning-to-agglomerate-superpixel-hierarchies.

22. Vazquez-Reina, A. et al. Segmentation fusion for connectomics. Proc. IEEE Int. Conf. on Comput. Vis. 177–184 (2011). DOI 10.1109/ICCV.2011.6126240.

23. Liu, T., Jurrus, E., Seyedhosseini, M., Ellisman, M. & Tasdizen, T. Watershed merge tree classification for electron microscopy image segmentation. In Pattern Recognition (ICPR), 2012 21st International Conference on, Icpr (2012). URL http://ieeexplore.ieee.org/xpls/abs_all.jsp?arnumber=6460090. DOI 10.1097/MPG.0b013e3181a15ae8.Screening.

24. Funke, J., Andres, B., Hamprecht, F. A., Cardona, A. & Cook, M. Efficient automatic 3D-reconstruction of branching neurons from em data. Proc. IEEE Comput. Soc. Conf. on Comput. Vis. Pattern Recognit. 1004–1011 (2012). DOI 10.1109/CVPR.2012.6247777.

25. Nunez-Iglesias, J., Kennedy, R., Parag, T., Shi, J. & Chklovskii, D. B. Machine learning of hierarchical clustering to segment 2D and 3D images. PLoS ONE 8 (2013). DOI 10.1371/journal.pone.0071715.

26. Parag, T., Chakraborty, A., Plaza, S. & Scheffer, L. A context-aware delayed agglomeration framework for electron microscopy segmentation. PLoS ONE 10, 1–19 (2015). DOI 10.1371/journal.pone.0125825.

27. Zaimi, A. et al. AxonDeepSeg: automatic axon and myelin segmentation from microscopy data using convolutional neural networks. Sci. Reports 1–11 (2017). URL http://arxiv.org/abs/1711.01004. DOI 10.1038/s41598-018-22181-4.

28. Dorkenwald, S. et al. Automated synaptic connectivity inference for volume electron microscopy. Nat. Methods 14, 435–442 (2017). URL http://dx.doi.org/10.1038/nmeth.4206. DOI 10.1038/nmeth.4206.

29. Kass, M., Witkin, A. & Terzopoulos, D. Snakes: Active contour models. Int. J. Comput. Vis. 1, 321–331 (1988). URL http://link.springer.com/10.1007/BF00133570. DOI 10.1007/BF00133570.

30. Jurrus, E. et al. Axon tracking in serial block-face scanning electron microscopy. Med. Image Analysis 13, 180–188 (2009). URL http://dx.doi.org/10.1016/j.media.2008.05.002. DOI 10.1016/j.media.2008.05.002.

31. Adams, R. & Bischof, L. Seeded region growing. IEEE Transactions on Pattern Analysis Mach. Intell. 16, 641–647 (1994).

32. Lucchi, A., Smith, K., Achanta, R., Knott, G. & Fua, P. Supervoxel-based segmentation of mitochondria in em image stacks with learned shape features. IEEE Transactions on Med. Imaging 31, 474–486 (2012). DOI 10.1109/TMI.2011.2171705.

33. West, K. L., Kelm, N. D., Carson, R. P. & Does, M. D. A revised model for estimating g-ratio from MRI. NeuroImage 125, 1155–1158 (2016). URL http://dx.doi.org/10.1016/j.neuroimage.2015.08.017. DOI 10.1016/j.neuroimage.2015.08.017.

34. Wang, L., Dong, J., Cull, G., Fortune, B. & Cioffi, G. A. Varicosities of Intraretinal Ganglion Cell Axons in Human and Nonhuman Primates. Investig. Ophthalmol. & Vis. Sci. 44, 2–9 (2003). DOI 10.1167/iovs.02-0333.

35. McDonald, J. H. Handbook of Biological Statistics. Sparky House Publ. 291 (2009). DOI 10.1017/CBO9781107415324.004.

36. Rushton, W. A. H. A theory of the effects of fiber size in medullated nerve. The J. Physiol. 101–122 (1951).

37. Stikov, N. et al. In vivo histology of the myelin g-ratio with magnetic resonance imaging. NeuroImage 118, 397–405 (2015). URL http://dx.doi.org/10.1016/j.neuroimage.2015.05.023. DOI 10.1016/j.neuroimage.2015.05.023.

38. Barazany, D., Basser, P. J. & Assaf, Y. In vivo measurement of axon diameter distribution in the corpus callosum of rat brain. Brain 132, 1210–1220 (2009). DOI 10.1093/brain/awp042.

39. Liewald, D., Miller, R., Logothetis, N., Wagner, H. J. & Schüz, A. Distribution of axon diameters in cortical white matter: an electron-microscopic study on three human brains and a macaque. Biol. Cybern. 108, 541–557 (2014). DOI 10.1007/s00422-014-0626-2.

40. Stikov, N. et al. Quantitative analysis of the myelin g-ratio from electron microscopy images of the macaque corpus callosum. Data Brief 4, 368–373 (2015). DOI 10.1016/j.dib.2015.05.019.

41. Greenberg, M. M., Leitao, C., Trogadis, J. & Stevens, J. K. Irregular geometries in normal unmyelinated axons: A 3D serial EM analysis. J. Neurocytol. 19, 978–988 (1990). URL http://link.springer.com/10.1007/BF01186825. DOI 10.1007/BF01186825.

42. Shepherd, G. M. G., Raastad, M. & Andersen, P. General and variable features of varicosity spacing along unmyelinated axons in the hippocampus and cerebellum. Proc. Natl. Acad. Sci. 99, 6340–6345 (2002). DOI 10.1073/pnas.052151299.

43. Kamiya, K. et al. Diffusion imaging of reversible and irreversible microstructural changes within the corticospinal tract in idiopathic normal pressure hydrocephalus. NeuroImage: Clin. 14, 663–671 (2017). URL http://dx.doi.org/10.1016/j.nicl.2017.03.003. DOI 10.1016/j.nicl.2017.03.003.

44. Topgaard, D. Multidimensional diffusion MRI. J.Magn. Reson. 275, 98–113 (2017). URL http://dx.doi.org/10.1016/j.jmr.2016.12.007. DOI 10.1016/j.jmr.2016.12.007.

45. Chomiak, T. & Hu, B. What is the optimal value of the g-ratio for myelinated fibers in the rat CNS? A theoretical approach. PLoS ONE 4 (2009). DOI 10.1371/journal.pone.0007754.

46. Ju, H., Hines, M. L. & Yu, Y. Cable energy function of cortical axons. Sci. Reports 6, 1–13 (2016). URL http://dx.doi.org/10.1038/srep29686. DOI 10.1038/srep29686.

47. Novikov, D. S., Jensen, J. H., Helpern, J. A. & Fieremans, E. Revealing mesoscopic structural universality with diffusion. Proc. Natl. Acad. Sci. 111, 5088–5093 (2014). URL http://www.pnas.org/cgi/doi/10.1073/pnas.1316944111. DOI 10.1073/pnas.1316944111.

48. Fieremans, E. et al. In vivo observation and biophysical interpretation of time-dependent diffusion in human white matter. NeuroImage 129, 414–427 (2016). URL http://dx.doi.org/10.1016/j.neuroimage.2016.01.018. DOI 10.1016/j.neuroimage.2016.01.018.

49. Palombo, M., Alexander, D. C. & Zhang, H. A generative model of realistic brain cells with application to numerical simulation of the diffusion-weighted MR signal. NeuroImage 188, 391–402 (2019). URL https://doi.org/10.1016/j.neuroimage.2018.12.025. DOI 10.1016/j.neuroimage.2018.12.025.

50. Lee, H. et al. Along-axon diameter variation and axonal orientation dispersion revealed with 3D electron microscopy: implications for quantifying brain white matter microstructure with histology and diffusion MRI. Brain Struct. Funct. (2019). URL http://link.springer.com/10.1007/s00429-019-01844-6. DOI 10.1007/s00429-019-01844-6.

51. Hassouna, M. S. & Farag, A. A. Multistencils fast marching methods: A highly accurate solution to the Eikonal equation on Cartesian domains. IEEE Transactions on Pattern Analysis Mach. Intell. 29, 1563–1574 (2007). DOI 10.1109/TPAMI.2007.1154.

52. Kim, J. H. & Juraska, J. M. Sex difference in the development of axon number in the splenium of the rat corpus callosum from postnatal day 15 through 60. Dev. Brain Res. 102, 77–85 (1997). DOI 10.1016/S0165-3806(97)00080-1.

53. Wake, H. et al. Nonsynaptic junctions on myelinating glia promote preferential myelination of electrically active axons. Nat. Commun. 6 (2015). DOI 10.1038/ncomms8844.

54. Dowding, I. & Haufe, S. Powerful Statistical Inference for Nested Data Using Sufficient Summary Statistics. Front. Hum. Neurosci. 12 (2018). URL http://journal.frontiersin.org/article/10.3389/fnhum.2018.00103/full. DOI 10.3389/fnhum.2018.00103.

55. Armstrong, R. C., Mierzwa, A. J., Marion, C. M. & Sullivan, G. M. White matter involvement after TBI: Clues to axon and myelin repair capacity. Exp. Neurol. 275, 328–333 (2016). URL http://dx.doi.org/10.1016/j.expneurol.2015.02.011. DOI 10.1016/j.expneurol.2015.02.011.

56. Reeves, T. M., Phillips, L. L. & Povlishock, J. T. Myelinated and unmyelinated axons of the corpus callosum differ in vulnerability and functional recovery following traumatic brain injury. Exp. Neurol. 196, 126–137 (2005). DOI 10.1016/j.expneurol.2005.07.014.

57. Dikranian, K. et al. Mild traumatic brain injury to the infant mouse causes robust white matter axonal degeneration which precedes apoptotic death of cortical and thalamic neurons. Exp. Neurol. 211, 551–560 (2008). DOI 10.1016/j.expneurol.2008.03.012.

58. Johnson, V. E., Stewart, W. & Smith, D. H. Axonal pathology in traumatic brain injury. Exp. Neurol. 246, 35–43 (2013). URL http://dx.doi.org/10.1016/j.expneurol.2012.01.013. DOI 10.1016/j.expneurol.2012.01.013.

59. Rodriguez-Paez, A. C., Brunschwig, J. P. & Bramlett, H. M. Light and electron microscopic assessment of progressive atrophy following moderate traumatic brain injury in the rat. Acta Neuropathol. 109, 603–616 (2005). DOI 10.1007/s00401-005-1010-z.

60. Mierzwa, A. J., Marion, C. M., Sullivan, G. M., McDaniel, D. P. & Armstrong, R. C. Components of myelin damage and repair in the progression of white matter pathology after mild traumatic brain injury. J Neuropathol Exp Neurol 74, 218–232 (2015). URL http://www.ncbi.nlm.nih.gov/pubmed/25668562. DOI 10.1097/NEN.0000000000000165.

61. Virtanen, J., Uusitalo, H., Palkama, A. & Kaufman, H. The effect of fixation on corneal endothelial cell dimensions and morphology in scanning electron microscopy. Acta ophthalmologica 62, 577–85 (1984). URL http://www.ncbi.nlm.nih.gov/pubmed/6435388.

62. Kharatishvili, I., Nissinen, J. P., McIntosh, T. K. & Pitkänen, A. A model of posttraumatic epilepsy induced by lateral fluid-percussion brain injury in rats. Neurosci. 140, 685–697 (2006). DOI 10.1016/j.neuroscience.2006.03.012.

63. Deerinck, T. et al. Enhancing serial block-face scanning electron microscopy to enable high resolution 3-D nanohistology of cells and tissues. Microsc. Microanal. 16, 1138–1139 (2010). DOI 10.1017/S14319276100.

64. Sim, K. S., Thong, J. T. L. & Phang, J. C. H. Effect of shot noise and secondary emission noise in scanning electron microscope images. Scanning 26, 36–40 (2006). URL http://onlinelibrary.wiley.com/doi/10.1002/sca.4950260106/abstract%5Cnhttp://doi.wiley.com/10.1002/sca.4950260106. DOI 10.1002/sca.4950260106.

65. Maggioni, M., Katkovnik, V., Egiazarian, K. & Foi, A. Nonlocal transform-domain filter for volumetric data denoising and reconstruction. IEEE Transactions on Image Process. 22, 119–133 (2013). DOI 10.1109/TIP.2012.2210725.

66. Canny, J. A computational approach to edge detection. IEEE Transactions on Pattern Analysis Mach. Intell. PAMI-8, 679–698 (1986). DOI 10.1109/TPAMI.1986.4767851.

67. Rosenfeld, A. & Pfaltz, J. L. Sequential operations in digital picture processing. J. ACM 13, 471–494 (1966). URL http://portal.acm.org/citation.cfm?doid=321356.321357. DOI 10.1145/321356.321357.

68. Achanta, R. et al. Slic Superpixels Technical Report. EPFL Tech. Rep. 149300 (2010).

69. Blum, H. A transformation for extracting new descriptors of shape. Model. for perception speech visual form 19, 362–380 (1967). URL papers2://publication/uuid/33A7D570-B63C-4E43-996A-4DE15D8EE75F.

70. Van Uitert, R. & Bitter, I. Subvoxel precise skeletons of volumetric data based on fast marching methods. Med. physics 34, 627–638 (2007). DOI 10.1118/1.2409238.

71. Sethian, J. A. A fast marching level set method for monotonically advancing fronts. Proc. Natl. Acad. Sci. 93, 1591–1595 (1996). URL http://www.pnas.org/cgi/doi/10.1073/pnas.93.4.1591. DOI 10.1073/pnas.93.4.1591.

72. van Heekeren, R. J., Faas, F. G. a. & van Vliet, L. J. Finding the minimum-cost path without cutting corners. Image Analysis: 15th Scand. Conf. SCIA 2007, Aalborg, Denmark 263–272 (2007). URL http://link.springer.com/10.1007/978-3-540-73040-8_27. DOI 10.1007/978-3-540-73040-8_27.

73. Haralick, R. M. & Shapiro, L. G. Computer and Robot Vision (Addison-Wesley Longman Publishing Co., Inc., Boston, MA, USA, 1992), 1st edn.

74. Jaccard, P. Nouvelles researches sur la distribution florale. Bull. de la Société vaudoise des sciences naturelles 44, 223–270 (1908).

75. Dice, L. R. Measures of the amount of ecologic association between species. Ecol. 26, 297–302 (1945). URL http://doi.wiley.com/10.2307/1932409. DOI 10.2307/1932409.

76. Kuhn, H. W. The Hungarian method for the assignment problem. Nav. Res. Logist. Q. 2, 83–97 (1955). DOI https://doi.org/10.1002/nav.3800020109.

77. Hodneland, E. et al. A unified framework for automated 3-D segmentation of surface-stained living cells and a comprehensive segmentation evaluation. IEEE Transactions on Med. Imaging 28, 720–738 (2009). DOI 10.1109/TMI.2008.2011522.

